# BICD2 is a centriolar protein that controls mother-daughter centriole engagement and the licensing of duplication

**DOI:** 10.1101/2025.01.04.631313

**Authors:** Lídia Montes-Ruiz, Núria Gallisà-Suñé, Laura Regué, Lorena Fernández-de-Larrea, Joan Roig

## Abstract

Centrosomes and the centrioles at their core have key cellular roles that require their duplication exactly once during each cell cycle. Abnormal centriole numbers can contribute to the onset of different pathologies and thus different mechanisms have evolved to tightly control centriole duplication. Importantly, centriole pairs are barred from duplicating until they go through a not completely understood process of licensing that includes the physical separation of older (mother) and younger (daughter) centrioles. Here we report that the dynein adaptor BICD2 is a centriolar protein with a previously unknown dynein-independent role controlling mother-daughter centriole engagement. We show that a pool of BICD2 resides at the centrosome, surrounding mother centrioles close to the daughter centriole. Removal of BICD2 results in premature centriole disengagement in G2 and early M and in centriole amplification in both non-transformed and transformed cells. We characterize the molecular determinants of BICD2 centriolar localization and suggest that this localization, and thus the role of BICD2 at the centriole, is controlled by phosphorylation. Our findings reveal a novel function of BICD2, independent of its ability to interact with dynein, which is crucial for the regulation of centriole licensing and the centrosome duplication cycle.

## Introduction

Centrosomes are membraneless protein assemblies that play a key role organizing microtubules into higher-order structures such as the mitotic spindle, cilia and flagella. They are present in most animal cells and are additionally involved in regulating cell movement, shape and intracellular organization, in establishing cell polarity and in facilitating different aspects of intracellular signaling (Nigg & Holland, 2018).

Each centrosome contains one or two microtubule-based cylinders called centrioles surrounded by pericentriolar material (PCM). The PCM is arranged in layers of coiled-coil rich proteins that surround the oldest centriole. It recruits different cellular components including microtubule nucleation complexes and during the G2 and M phases of the cell cycle it expands as a matrix-like assembly. This enhances its ability to organize microtubules and assemble a strong microtubule aster for spindle formation. At the core of centrosomes, centrioles are structurally well-defined entities, precisely formed around a cylindrical scaffold of microtubule triplets arranged in nine-fold symmetry and specific polarity. Different proteins localize at distinct regions of this microtubule cylinder, defining the centriole proximal, central and distal ends, the first and the last corresponding to the microtubule minus and plus ends, respectively (LeGuennec *et al*, 2021; Laporte *et al*, 2024). To form a single cilium in G1/G0 and two spindle poles in mitosis, the number of centrioles (and thus of centrosomes) needs to be tightly controlled. For this, two different regulatory systems ensure that centrioles are duplicated only once during the S phase of each cell cycle. The first system involves PLK4, a protein kinase responsible for regulating the initiation of centriole duplication, and ensuring that only one “daughter” (new) centriole is assembled for each “mother” (preexisting) centriole. Briefly, active PLK4 accumulates in a single dot-like structure at the centriole wall alongside SAS-6, one of the main organizers of forming centrioles. Phosphorylation of SAS-6 and other proteins then leads to the growth of a procentriole around a nine-fold symmetric structure known as the cartwheel (Arquint & Nigg, 2016; Gönczy & Hatzopoulos, 2019).

The described process can only happen if the two original centrioles that the cell receives in G1 are separated and thus “licensed” to become proficient mother centrioles. Control of licensing (Tsou & Stearns, 2006a) is thus the second regulatory system that ensures that a single centriole is produced for each preexisting centriole once and only once each cell division cycle. Until G1 mother/daughter centriole pairs (called diplosomes) are tightly associated (“engaged”) and arranged perpendicularly in a configuration that is not competent for centriole duplication (Wong & Stearns, 2003). The presence of a daughter associated with the mother centriole inhibits the formation of additional centriole daughters (Loncarek *et al*, 2008). During late mitosis, centrioles “disengage”, losing their associated configuration through a mechanism that involves the anaphase-promoting complex and the protease separase (Tsou & Stearns, 2006b). After disengagement, the older centriole is ready to duplicate while the youngest one still needs to go through a maturing process called centriole-to-centrosome conversion in which it will acquire different new proteins including those necessary for PLK4 recruitment (importantly CEP192 and CEP152), ultimately resulting in the organization of its own PCM (Wang *et al*, 2011; Kim *et al*, 2013; Sonnen *et al*, 2013; Izquierdo *et al*, 2014; Tsuchiya *et al*, 2016; Fu *et al*, 2016). Around the same time in human cells younger centrioles will lose their inner scaffold, the cartwheel, a step that may also be important for licensing (Fong *et al*, 2014; Kim *et al*, 2016; Huang *et al*, 2022).

Both disengagement and centriole-to-centrosome conversion are processes regulated by the protein kinase PLK1 (Tsou *et al*, 2009; Loncarek *et al*, 2010; Wang *et al*, 2011; Shukla *et al*, 2015), possibly through the direct phosphorylation of a number of centriolar proteins in conjunction with the main mitotic regulator CDK1. During disengagement PLK1 acts independently of separase and, in fact, the kinase alone is able to induce the separation of mother and daughter centrioles (Tsou *et al*, 2009; Hatano & Sluder, 2012; Shukla *et al*, 2015).

The nature of the proteins that keep daughter centrioles engaged with their mothers, as well as how separase and mitotic phosphorylation regulate them has been a matter of controversy. Separase localizes to the centrosomes until anaphase (Chestukhin *et al*, 2003) and its proteolytic activity has been detected at the organelles (Agircan & Schiebel, 2014) suggesting that its main substrate, the ring-like protein complex cohesin, might also be located at centrosomes and have a direct role controlling centriole engagement, similarly to its role in regulating DNA sister chromatid cohesion. The hypothesis that cohesin cleavage drives centriole disengagement has received support from different studies (reviewed in (Simmons Kovacs & Haase, 2010)). For example, ectopic cohesin ring opening in *Xenopus* egg extracts can induce centriole disengagement, while this is blocked by non-cleavable cohesin subunits (Schöckel *et al*, 2011). However, conflicting evidence exists. Tsou and collaborators (Tsou *et al*, 2009) reported contrasting results and a study in *Drosophila* convincingly demonstrates that cohesin ring opening is insufficient for centriole disengagement (Oliveira & Nasmyth, 2013). Moreover, in *C. elegans,* cohesin cleavage is only required in meiosis II and not during mitotic cycles (Cabral *et al*, 2013). Therefore, the role of cohesin in regulating engagement remains a subject of debate. It may be that the complex is only relevant for the control of (dis)engagement in some specific conditions (i.e. in meiotic cells). Importantly, cohesin has not been specifically observed at mother-daughter centriole contact zones.

Thus, other possible separase substrates have been sought after to account for the role of the protease at the centrioles. The most convincing candidate to date is pericentrin (known also as kendrin in its alternatively spliced form), a more than 3000 residue-long, coiled-coil rich protein that is a key scaffolding component of the PCM. Pericentrin is a target of separase, and its cleavage has been proposed to contribute to disengagement. Moreover, expression of non-cleavable forms of the protein suppresses normal disengagement (Matsuo *et al*, 2012; Lee & Rhee, 2012; Kim *et al*, 2019). Pericentrin may be assisted in keeping centriole engagement by other PCM proteins such as CDK5RAP2 (Pagan *et al*, 2015) and enticingly has been shown to depend on PLK1 phosphorylation to be an efficient separase target (Kim *et al*, 2015), although this is discussed (Agircan & Schiebel, 2014). It is anchored to the mother centriole through CEP57, which is also in charge of assembling a toroid around the proximal region that contains CEP152, crucial for PLK4 and PCM recruitment (Watanabe *et al*, 2019; Ito *et al*, 2021). How pericentrin may connect mother and daughter centrioles is not completely understood as no specific interaction of the protein with daughter centriole components has been identified. Given the critical roles of pericentrin at the PCM, it is also possible that the entire pericentriolar material contributes to maintaining centriole engagement. It is not clear how the cleavage of pericentrin would be functionally important in this case. PCM disassembly, helped by astral microtubules pulling forces plus cytoskeletal dynamics may suffice to disengage centrioles without the need to cleave it, as observed in *C.elegans* (Cabral *et al*, 2013).

We present here results that show that the dynein adaptor BICD2, a coiled-coil rich protein with roles in vesicle transport in interphase and in the recruitment of the dynein motor complex to the nuclear envelope during G2 and early mitosis (Hoogenraad & Akhmanova, 2016), is a centrosomal protein involved in keeping daughter and mother centrioles engaged. Our findings demonstrate that disengagement results from the removal of BICD2 in cells that have normal amounts of pericentrin at the centrosomes, thus arguing against the need of pericentrin cleavage or PCM disassembly for centriole disengagement. Moreover, we present data indicating that BICD2 roles at the centrioles depend on the ability of its C-terminal region to assemble around them and are independent of its function as a dynein adaptor/activator. We finally suggest a mechanism for the removal of BICD2 from centrioles during G2 and mitosis, i.e. its C-terminal phosphorylation by mitotic kinases, and discuss the possibility that other regulatory mechanisms exist that may completely remove the protein during anaphase.

Defective mother-daughter engagement can lead to premature disengagement and centriole amplification, something that we observed in both non-transformed and transformed cells with abnormal levels of BICD2. Centriole amplification results in different pathological conditions and is associated with the transformed phenotype of cancer cells (Goundiam & Basto, 2021). Our findings thus not only reveal BICD2 as a crucial factor in the centriole duplication cycle, but lay the groundwork for future studies exploring the potential link between BICD2 misfunction and human disease.

## Results

During our work studying the regulation of BICD2 by phosphorylation (Gallisà-Suñé *et al*, 2023), we noticed that a significant pool of the protein colocalized with different centrosomal markers. This is exemplified in Figure 1A, showing non-transformed retinal pigment epithelial cells (hTERT RPE-1 cells, henceforth RPE-1 cells) stained for BICD2 and the centriolar protein CEP152.

**Figure 1.**
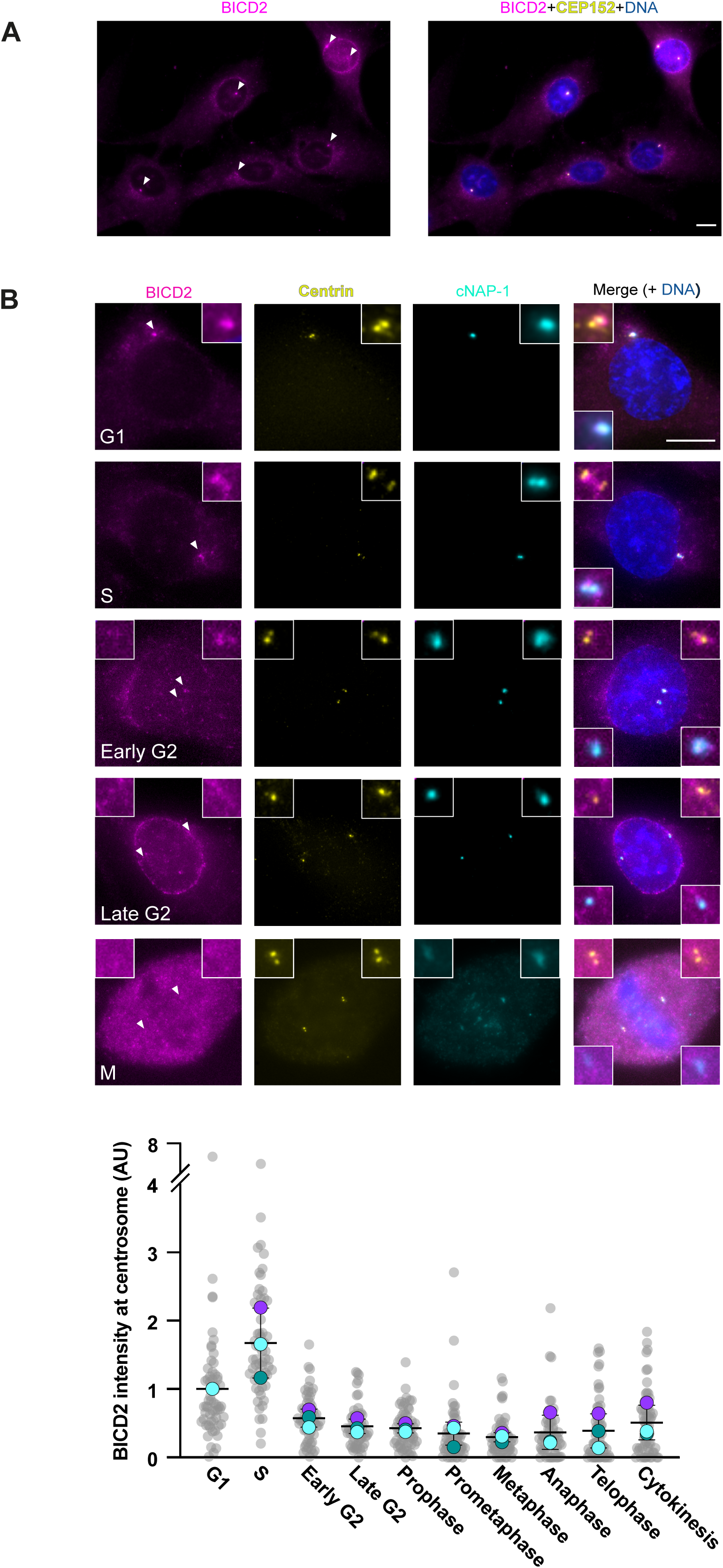
A pool of BICD2 localizes to the centrosome in a cell cycle-regulated manner. **A.** Immunofluorescence images of RPE-1 cells stained for BICD2 (GT10811 antibody), centrosomes (CEP152) and DNA. Arrowheads point to the centrosomes. Scale bar 10 µm. **B.** *Top*, examples of cells in different cell cycle phases stained for BICD2 (GT10811 antibody), centrin, the mother centriole marker C-NAP1 plus DNA. *Bottom*, quantification of the centrosomal intensity of the BICD2 signal in different phases of the cell cycle (n=3 biological replicates, 20 centrosomes per phase and replicate; individual replicate means plus mean ± SD of replicates are shown). Scale bar 10 µm.

Similar results were observed when co-staining with the centriolar markers centrin and C-NAP1/CEP250 (Figure 1B). BICD2 amounts at the centrosome changed with the different phases of the cell cycle (Figure 1B; see also Supplementary Figure 1). They were high in G1 and S, when, as previously described (Hoogenraad *et al*, 2001), a pool of BICD2 also localized to the Golgi apparatus (Supplementary Figure 1). During G2, when in response to different regulatory inputs the bulk of BICD2 interacts with the nucleoporin RanBP2 and localizes to the nuclear envelope (Splinter *et al*, 2010), we observed that the amount of centrosomal BICD2 started to diminish (being in late G2, when the centrosomes are completely separated across the nucleus, 45±11% of the value observed in G1). The amount of centrosomal BICD2 reached a minimum in mid-mitosis (29±7% of the value in G1 in metaphase), a phase when in some cells the protein was barely detectable at centrosomes. Centrosomal BICD2 amounts started to increase after mitosis, reaching maximum levels again in G1 and then S phases.

The use of different anti-BICD2 antibodies confirmed the specificity of our observations (Supplementary Figure 2A), which were further supported by our results in BICD2 knockout cells (see Figure 5B below). We also detected BICD2 at centrosomes in other cell lines, including U2OS osteosarcoma cells, where protein levels at centrosomes oscillated similarly to those in RPE-1 cells (Supplementary Figure 2B).

Importantly, depolymerization of the cell microtubules using cold treatment did not significatively change the amount of BICD2 at centrosomes (Supplementary Figure 3A), indicating that the observed localization is not the result of the accumulation of the adapter at the minus ends of pericentrosomal microtubules (i.e. as part of dynein motor complexes). Furthermore, the result suggested that the centrosomal localization of BICD2 does not depend (at least for short periods of time) on the presence of the microtubule cytoskeleton.

Comparisons of the staining patterns of BICD2 to those observed for different markers such as centrin, C-NAP1 or CEP152 (e.g. in Figure 1 and Supplementary Figure 1) strongly suggested that BICD2 localizes to the centrioles, and specifically to the mother centriole, and not to the PCM. In fact the PCM is most abundant in G2 and mitosis, after centrosome maturation, precisely when centrosomal BICD2 is minimal. Furthermore, we observed that BICD2 localization did not overlap with that of pericentrin, one of the key components of the PCM in human cells (Supplementary Figure 3B). The bulk of BICD2 neither seemed to localize to centriolar satellites, as determined by co-staining with the satellite component PCM-1 (Supplementary Figure 3B).

To better describe BICD2 localization at the centriole we used super-resolution microscopy. Our results with RPE-1 cells using three-dimensional structured illumination microscopy (3D-SIM) are shown in Figure 2, Figure 3 and Supplementary Figures 4 and 5. In cells with two non-separated diplosomes (i.e., in S phase, showing the higher amounts of centriolar BICD2), BICD2 accumulated around the mother centrioles (Figure 2), partially colocalizing with CEP152, that forms a ring at the proximal half of mature centrioles. The localization pattern of BICD2 was more variable than that of CEP152, not always forming a closed ring and in many cases consisting of a discontinuous ring-shaped structure around the centriole. BICD2 also tended to occupy a wider region than CEP152. This was more patently observed with one of the two different antibodies that we used, GT10811, that tended to produce staining patterns that surrounded the centrioles externally to CEP152 and also to the signal of the ab117818 antibody (compare Figure 2A and 2B and also see Supplementary Figure 4A). The results suggest that the different epitopes recognized by the two antibodies lay far apart and that BICD2 may have an at least partially extended conformation at the centriole, although this will need to be studied further. The longitudinal location of BICD2 at the mother centriole was coincident with the position of the daughter centriole, around the base of which, identified by the cartwheel component SAS-6, we noted that BICD2 tended to accumulate. This was supported by data with the daughter centriole marker centrobin (Supplementary Figure 4B). As expected from the previous results, we observed that BICD2 localized between CEP164, a centriolar distal appendage component, and C-NAP1, a component of the intercentriolar link that accumulates at the proximal end of mature centrioles (Supplementary Figure 4C). Finally, super-resolution images of cells stained with BICD2 and pericentrin, confirmed that the two proteins did not colocalize in interphase and that BICD2 almost disappeared from the PCM of mitotic cells, when the amount of pericentrin at the centrosome reaches its maximum (Supplementary Figure 4D).

**Figure 2.**
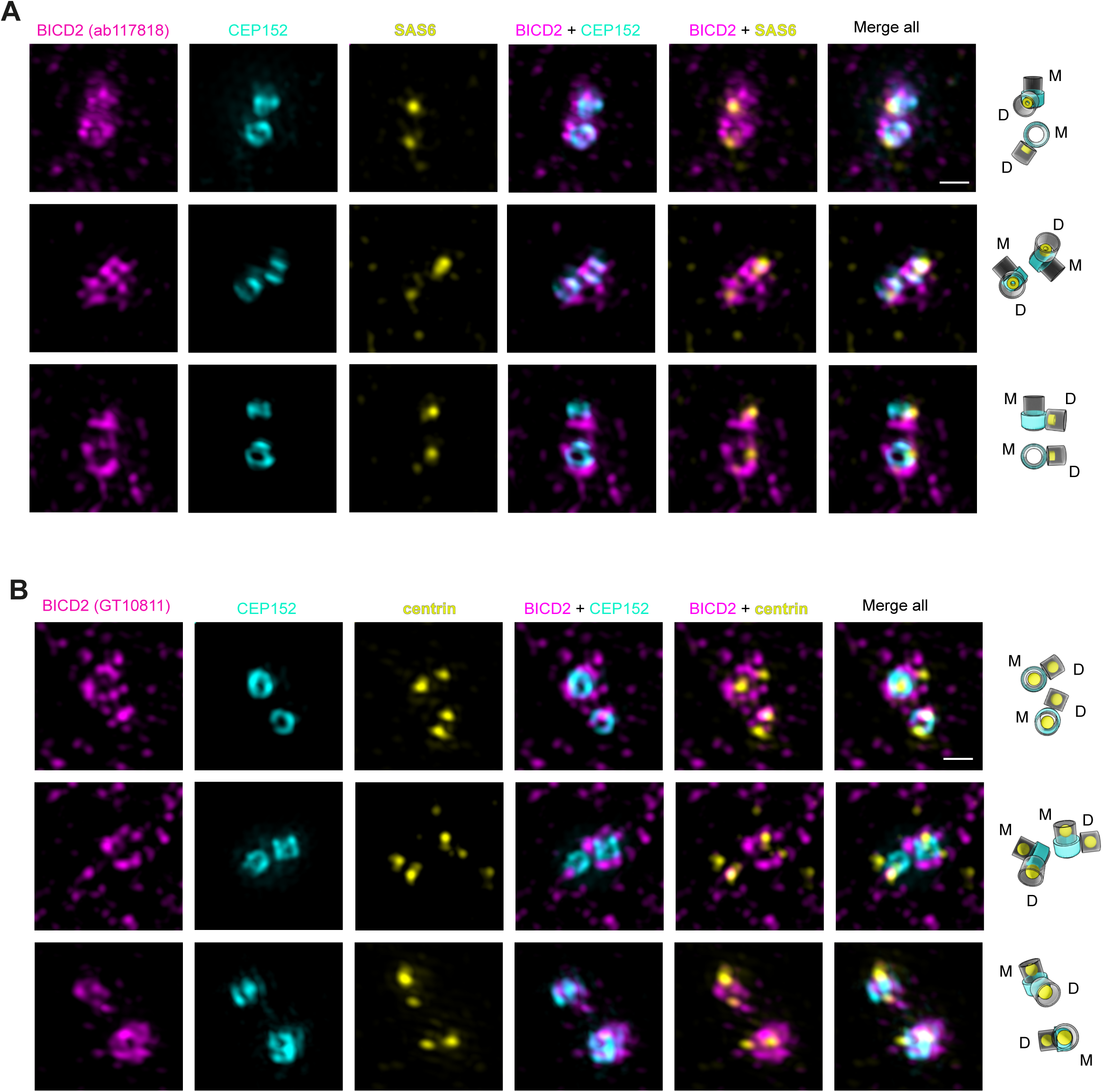
BICD2 localizes on the mother centriole at the level of the daughter centriole. **A.** 3D-Structured illumination microscopy (SIM) images of centrosomes of RPE-1 cells in S phase (two diplosomes), stained for BICD2 (ab117818 antibody), CEP152 and SAS-6. CEP152 identifies the mother centriole (M) while SAS-6 labels the daughter centriole (D). Maximum projections of z-stacks are shown. Scale bar 0.5 µm. **B.** As in A, cells stained for BICD2 (GT10811 antibody), CEP152 and centrin.

**Figure 3.**
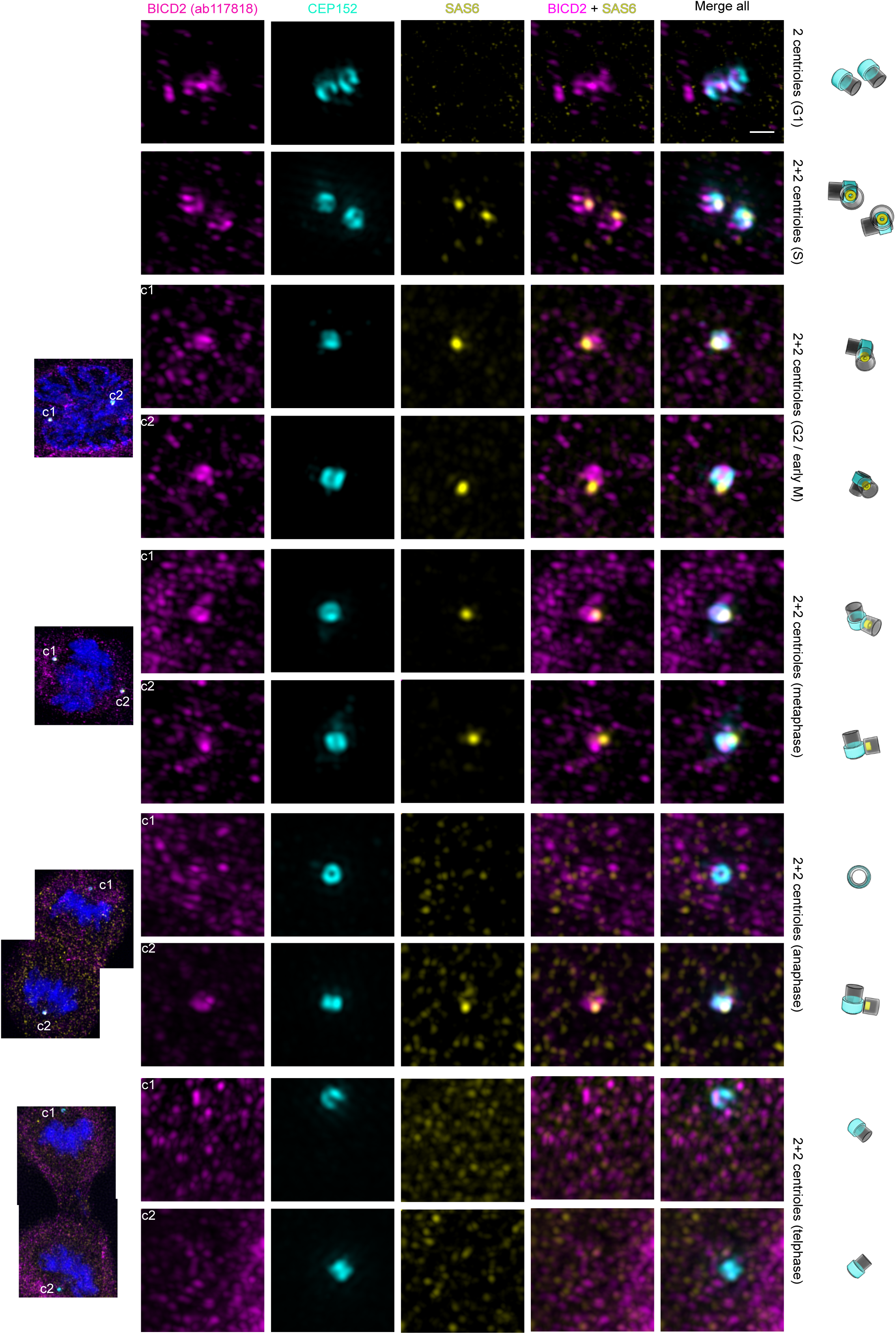
Centriolar BICD2 during the cell cycle. 3D-SIM images of centrosomes of RPE-1 cells in the indicated phases of the cell cycle stained for BICD2 (ab117818 antibody), CEP152 and SAS-6. CEP152 identifies the mother centriole while SAS-6 labels the daughter centriole. Maximum projections of z-stacks are shown. An overview image of the cell including DNA staining is additionally shown for mitotic cells. Scale bar 0.5 µm.

We next used super-resolution microscopy to study in detail how the distribution of BICD2 at the centriole changed during its duplication cycle. In RPE-1 cells with a single pair of centrioles (G1 phase), BICD2 showed either a disorganized pattern or a ring-like shape around CEP152-positive centrioles (Figure 3 and Supplementary Figure 5). Cells in G1 can have one or two CEP152-positive centrioles, as daughter centrioles gain CEP152, among other proteins, while being converted to mature mother centrioles during this phase. We observed that in G1 cells with two CEP152-positive centrioles one of them often showed reduced or negligible amounts of BICD2 (Supplementary Figure 5). We interpret our observations as showing that in G1 BICD2 accumulates first to the older mother centriole and then to the younger centriole, after this has acquired CEP152.

In S phase (four centrioles organized in two diplosomes) both mother centrioles were positive for BICD2, and as mentioned before the protein tended to embrace the mother centriole at the level of the daughter centriole, often accumulating close to its base and the cartwheel, visualized by SAS-6 (Figure 3 and Supplementary Figure 5).

The amount of centriolar BICD2 diminished in G2 and early M cells, with two separated centrosomes, but remained visible in both diplosomes, associated with the SAS-6 foci (Figure 3). Around anaphase BICD2 was not detectable in some of the diplosomes, while some others retained some protein (again, close by the SAS-6 foci, that at this point was disappearing from the centrioles as the result of the disassembly of the cartwheel). Later in mitosis, around telophase and during cytokinesis most of the diplosomes were negative for BICD2 (Figure 3 and Supplementary Figure 5). We obtained similar results in U2OS cells using both 3D-SIM (Supplementary Figure 6) and image scanning microscopy/Airyscan (Supplementary Figure 7).

We next investigated which part of BICD2 was responsible for its observed centriolar localization. For this we expressed different forms of GFP-tagged BICD2 in RPE-1 cells. Figure 4A shows that full-length BICD2 fused to GFP is able to localize to the centrosomes in RPE-1 cells, although this localization was sometimes obscured by the amount of protein accumulated around the organelle, possibly as a result of the ability of full-length BICD2 to activate dynein and concentrate at the microtubules minus ends. Depolymerizing microtubules by incubating cells on ice before fixation allowed us to better observe the centrosomal localization of GFP-BICD2. GFP-BICD2 [1-575] containing the N-terminal dynein-binding part of the adaptor did not accumulate significantly at centrosomes in RPE-1 cells. Importantly, different C-terminal fragments of BICD2 clearly localized to the centrosome, with a minimal fragment comprising residues 658-820, comprising the last coiled-coil segment of the protein (CC4), strongly localizing to the organelle. Shorter fragments did not localize so clearly, and a short polypeptide comprising the last residues after the CC4 region, BICD2 [802-820], was not observed at the centrosome. Thus, we conclude that BICD2 localizes to the centrosome through its C-terminal region. Our results also indicated that the centrosomal localization of BICD2 is dynein-independent, as the motor complex interacts with the N-terminal region of the adaptor and thus C-terminal fragments must localize independently of the motor (e.g. (Hoogenraad *et al*, 2003; McKenney *et al*, 2014; Schlager *et al*, 2014; Lee *et al*, 2018)).

**Figure 4.**
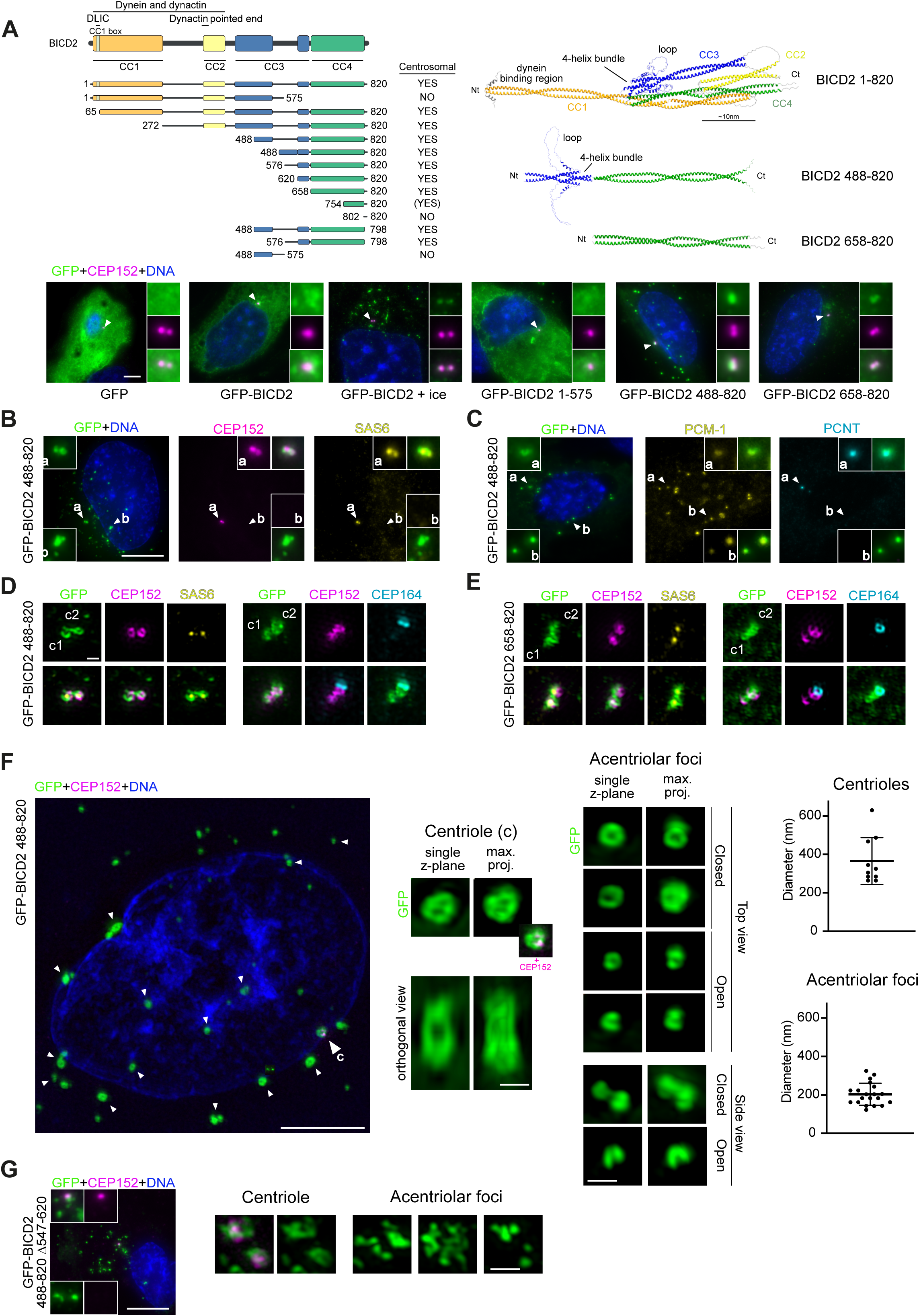
BICD2 localizes to the centriole through its C-terminus, which is able to assemble in toroidal structures. **A.** *Left*, schematic representation of BICD2, noting its coiled coil (CC) domains as predicted by AlphaFold and the regions interacting with the dynein/dinactin motor complex. The different fragments tested for centrosomal localization are depicted below. Expression of the different GFP-fusion proteins is shown in Supplementary Figure 8A. *Right*, AlphaFold structure prediction for the full-length protein (residues 1-820) plus the two C-terminal fragments studied in detail below (residues 488-820 and 658-820). Note that all residue numbers refer to the mouse sequence. Human BICD2 was predicted to have a fundamentally identical structure. *Bottom*, example immunofluorescence images of RPE-1 cells expressing the indicated GFP-fusion polypeptides and stained for GFP, CEP152 as a centrosome marker and DNA. *ice* indicates that the cells were incubated over ice 30 minutes before fixation in order to depolymerize microtubules. Arrowheads point to the centrosomes. Scale bar 10 µm **B.** Immunofluorescence images of RPE-1 cells expressing GFP-BICD2[488-820] and stained for GFP, CEP152, SAS-6 and DNA. Arrowheads point to the centrosomes (*a*) or to non-centrosomal GFP foci (*b*). Scale bar 10 µm. **C.** As in B, cells stained for GFP, PCM1, pericentrin (PCNT) and DNA. **D.** 3D-SIM images of centrosomes (labeled *c1* and *c2*) of RPE-1 cells expressing GFP-BICD2[488-820] stained for GFP, CEP152 and SAS-6 (left panels) or GFP, CEP152 and CEP164 (right panels). Maximum projections of z-stacks are shown. Scale bar 0.5 µm. **E.** As in *D*, cells expressing GFP-BICD2[658-820]. **F.** *Left*, 3D-SIM images of an RPE-1 cell expressing GFP-BICD2[488-820] stained for GFP, CEP152 as a centrosomal marker and DNA. A maximum projection of the z-stack is shown. Scale bar 5 µm. *Center*, single z-planes and maximum projections of centrioles and example acentriolar GFP foci. An orthogonal view of the z-stack is also shown for the centriole. Scale bar 0.5 µm. *Right*, diameter of the centriolar and acentriolar toroidal GFP structures observed (n=10 centrioles, n=20 acentriolar foci; mean ± SD is shown). **G.** *Left*, immunofluorescence images of a cell expressing a GFP-BICD2[488-820] form lacking the disorganized loop interrupting the coiled-coil region (GFP-BICD2[488-820, Δ547-620]) stained for GFP, CEP152 as a centrosomal marker, and DNA. Scale bar 10 µm. *Right*, example 3D-SIM images of centrioles and acentriolar GFP foci of similar cells. Scale bar 0.5 µm. In all cases maximum projections of the z-stack are shown. Expression of the different GFP-fusion proteins is shown in Supplementary Figure 8A.

We further characterized the subcellular localization of two C-terminal forms of BICD2, respectively comprising residues 488-820 and 658-820. Both polypeptides were predicted by AlphaFold (Abramson *et al*, 2024) to form extended coiled-coil dimers (Figure 4A). Interestingly, BICD2 [488-820], that contains a short disorganized region that interrupted the CC3 coiled-coil region (residues 547-622), was predicted to form a four-helix bundle through an intramolecular interaction that results in a linear dimeric structure from which two disorganized loops protrude. As shown in Figure 4A, both GFP-BICD2 [488-820] and GFP-BICD2 [658-820] localized to the centrosomes, but while the shorter GFP-BICD2 [658-820] accumulated almost exclusively at the organelle, GFP-BICD2 [488-820] additionally formed a number of cytoplasmic foci. These extracentrosomal foci did not contain centrosomal markers such as CEP152, SAS-6 (Figure 4B) or pericentrin (Figure 4C), although, strikingly, most if not all of GFP-BICD2 [488-820] foci colocalized with the centriolar satellite marker PCM-1 (Figure 4C). The presence of BICD2 at the centriolar satellites has been suggested previously (Quarantotti *et al*, 2019); we ignore the reason why in the conditions used in this study only C-terminal fragments of BICD2 but not the full-length protein (see Supplementary Figure 3B) colocalize with these pericentriolar structures. Using 3D-SIM we observed that GFP-BICD2 [488-820] and GFP-BICD2 [658-820] accumulated around the centrioles, forming structures that in the case of GFP-BICD2 [488-820] were more frequently ring-like shaped. In both cases the polypeptides partially colocalized with CEP152 and engulfed SAS-6 foci (Figures 4D and 4E). We further study the structures formed by GFP-BICD2 [488-820]. Around the centriole this polypeptide formed rings or cylinders of an average diameter of ∼400 nm (Figure 4F). Strikingly, most acentriolar foci of GFP-BICD2 [488-820] also showed a ring-like morphology, some closed some open, with a smaller diameter of ∼200nm. Moreover, a form of the polypeptide that lacked the disordered loop in CC3, GFP-BICD2 [488-820 Δ547-620], although centrosomal, failed to form the observed ring-like structures in the cytoplasm (Figure 4G). In view of our results, we propose that the C-terminal part of BICD2 is not only able to localize to the centriole, but also has the intrinsic tendency to associate into a ring-like structure, a property that depends on its unstructured loop in the CC3 region and may facilitate BICD2 specific centriolar disposition.

The spatiotemporal pattern of BICD2 localization, with the accumulation of the protein at the proximal part of the mother centrioles around the region where daughter centrioles are localized, as well as its disappearance in mid-late mitosis, led us to hypothesize that BICD2 might be involved in maintaining the engagement between the mother and daughter centrioles and in the regulation of centriole duplication licensing. To study this we produced two independent RPE-1 BICD2 knockout cell lines through the use of CRISPR-Cas9 genome editing, that we called NΔ4 and NΔ5 (Figure 5). Western blots confirmed that no BICD2 was expressed in the knockout cell lines, and DNA sequencing revealed the genetic changes responsible for this (Figure 5A). Importantly, the centrosomal BICD2 signal observed in wild type cells was conspicuously reduced in the knockout cell lines while that of other centrosomal markers such as pericentrin did not vary significatively (Figure 5B).

**Figure 5.**
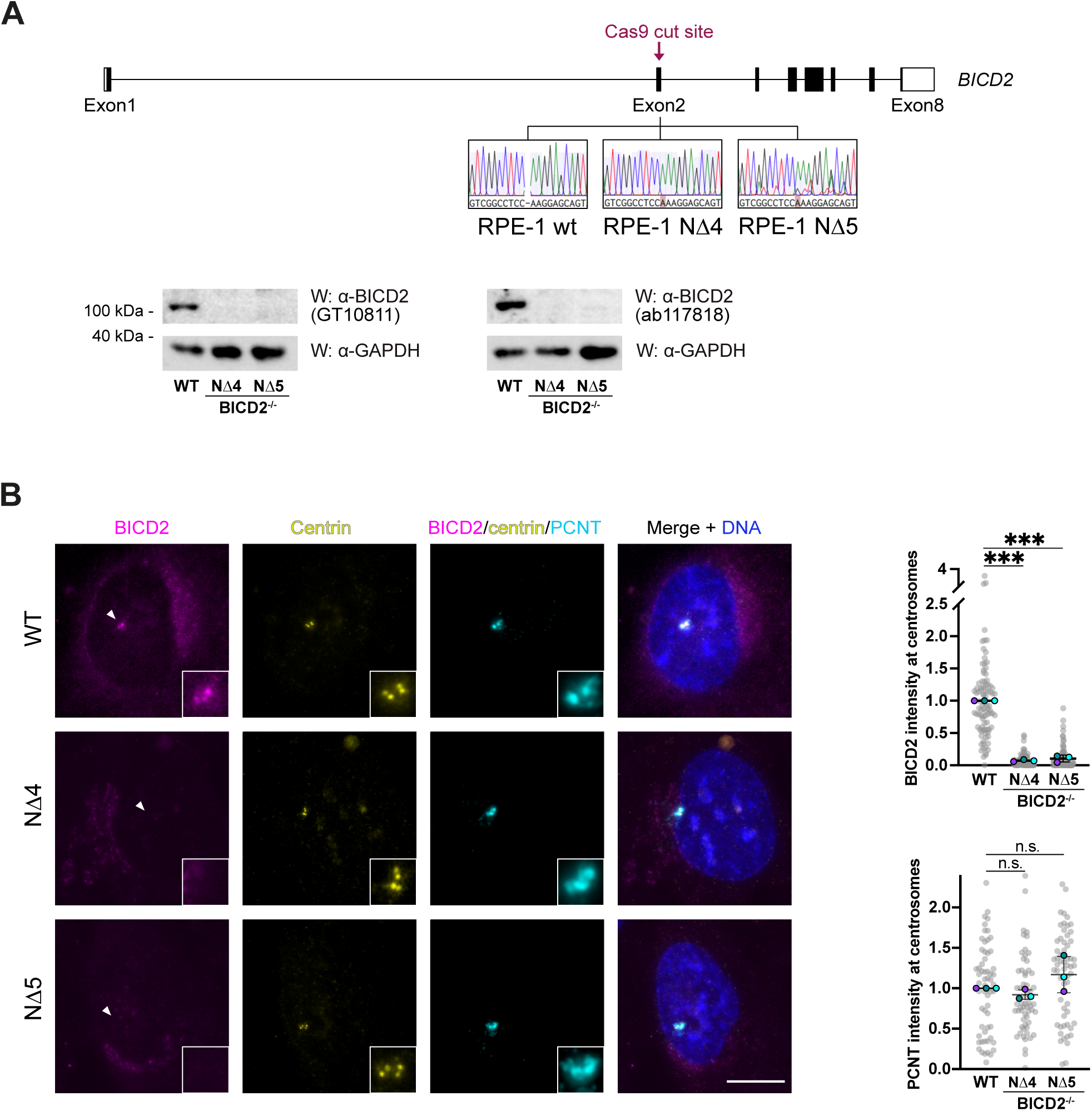
BICD2^-/-^ cells. **A.** BICD2^-/-^ cell lines were produced as described in Experimental Procedures. *Top*, schematic representation of the BICD2 gene (GeneID 23299) showing the engineered Cas9 cut site in exon 2 and the resulting base insertion in the two RPE-1 BICD2^-/-^ cell lines produced. The same DNA region is also shown for wild type cells. *Bottom*, western blots (*W*) of total cell extracts using different anti-BICD2 antibodies. GAPDH amounts are shown as loading controls. **B.** Example immunofluorescence images of control and BICD2^-/-^ cells stained for BICD2 (GT10811 antibody), centrin, pericentrin (PCNT) and DNA. Quantification of the amount of BICD2 (in G1/S cells, when maximal) and pericentrin (in mitotic cells, when maximal) at centrosomes is also shown (n=3 biological replicates, 30 centrosomes per replicate; values normalized to the mean values of control cells; individual replicate means plus mean ± SD of replicates are shown; statistical significance analyzed using one-way ANOVA with post hoc analysis). Scale bar 10 µm

In order to assess the importance of BICD2 for mother-daughter centriole engagement, we measured the distance between centrioles for each centriole pair (i.e. diplosome) in the BICD2^-/-^ cell lines and their wilt type counterparts. We started by doing so in S phase, when centrioles have just duplicated and cells have 4 centrioles organized in two closely associated diplosomes, and did not observe any consequences of the lack of BICD2 (Figure 6A). In RPE-1 cells intercentriolar distances, as measured using centrin as a centriole marker, were in this phase similar to those described in HeLa cells (Shukla *et al*, 2015), i.e. 0.4-0.5µm. We next measured intercentriolar distances in G2, observing that BICD2^-/-^ cells showed slightly, but significantly, bigger distances than wild type cells (0.54±0.04µm for both knockout cells lines vs 0.48±0.01µm in wild type cells; Figure 6B). This effect was more pronounced in cells in early mitosis (up to metaphase, before licensing usually takes place, Figure 6C), with wild type cells showing a mean intercentriolar distance of 0.43±0.04 µm, and BICD2^-/-^ cell lines of 0.59±0.04 and 0.66±0.07µm. Notably, in the knockout cells we observed cells with centrioles more than 1 µm apart, while in the wild type cells no distances bigger than 0.7µm were observed. This effect was magnified in cells arrested in prometaphase by the use of the Eg5/KIF11 inhibitor STLC (DeBonis *et al*, 2004). In this case wild type RPE-1 cells showed a mean intercentriolar distance of 0.96±0.14µm while the two BICD2^-/-^ cell lines showed distance that were significatively higher, 1.36±0.15µm and 1.46±0.19µm (Figure 6D).

**Figure 6.**
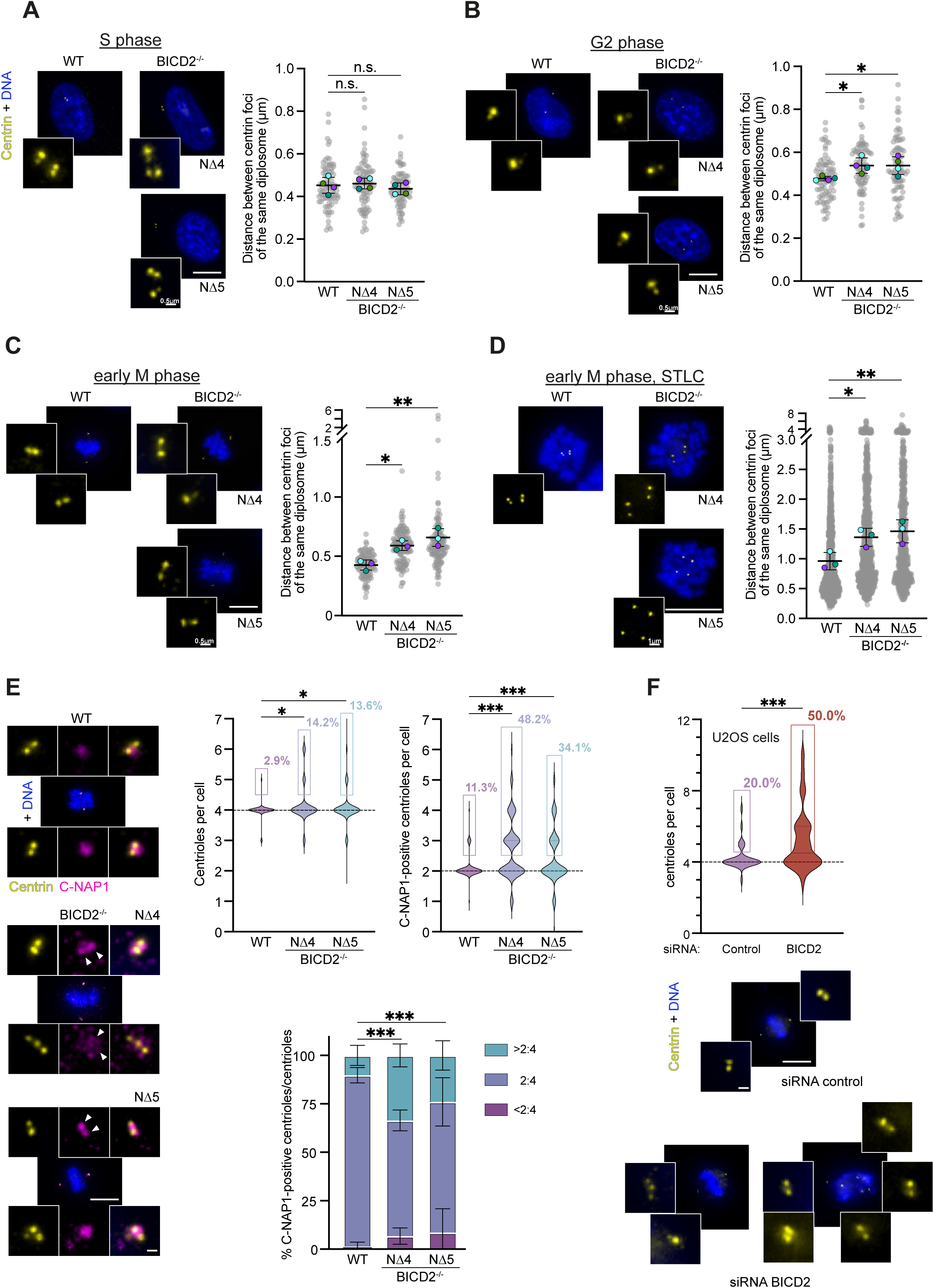
Abnormal BICD2 amounts result in increased intercentriolar distances in G2 and M and in centriole amplification. **A.** Distance between centrin foci of the same diplosome (defined as the cellular pair of centrioles with the smallest intercentriolar distance) in wild type and BICD2^-/-^ RPE-1 cells in S phase, i.e. with two closely associated diplosomes (n=4 biological replicates, 20 diplosomes per replicate; individual replicate means plus mean ± SD of replicates are shown and statistical significance is analyzed using one-way ANOVA with post hoc analysis). **B.** As in *A*, RPE-1 cells in G2, i.e. with two clearly separated diplosomes and uncondensed DNA (n=4 biological replicates, 20 diplosomes per replicate). **C.** As in *A*, RPE-1 cells in early mitosis (prophase, prometaphase or metaphase; two separated diplosomes and DNA condensed, phase assigned by observation of the chromosomes; n=3 biological replicates, 20-50 diplosomes per replicate). **D.** As in *A*, RPE-1 cells treated overnight with STLC and thus arrested in prometaphase (n=3 biological replicates, ∼200 diplosomes per replicate). **E.** *Upper left*, number of centrioles (identified using centrin) per cell in wild type and BICD2^-/-^ mitotic RPE-1 cells (n=4 biological replicates, 20-30 cells per replicate; statistical significance analyzed using a Chi square test). The overall percentage of cells with >4 centrioles is additionally shown. *Upper right*, number of C-NAP1 positive centrioles per cell in early mitotic wild type and BICD2^-/-^ RPE-1 cells (n=3 biological replicates, 20-30 cells per replicate; statistical significance analyzed using a Chi square test). The overall percentage of cells with >2 C-NAP1 positive centrioles is shown. *Bottom*, percentage of cells with the indicated number of C-NAP1-positive centrioles/total number of centrioles ratios in wild type and BICD2^-/-^ mitotic RPE-1 cells (n=3 biological replicates, ∼30 cells per replicate; statistical significance analyzed using a Chi square test). **F.** U2OS cells were transfected with the indicated siRNAs as described in experimental methods and centriole numbers in mitotic cells were quantified using centrin as a marker (n=4 biological replicates, 10 cells per replicate; statistical significance analyzed using a Chi square test). RNAi efficiency was confirmed by western blot, see Supplementary Figure 8B. In all cases, immunofluorescence images of cells stained for centrin and DNA (A-D, F) or centrin, C-NAP1 and DNA (E) are shown. Scale bars correspond to 10 µm and 0.5 µm (insets).

The intercentriolar distances observed in mitotic BICD2^-/-^ RPE-1 cells are similar to those observed in anaphase HeLa cells and thus suggestive of disengagement (Shukla *et al*, 2015). Given that disengagement results in the licensing of duplication we could expect that BICD2^-/-^ cells would exhibit abnormal centriole amplification. To investigate this, we quantified numbers of centrioles in both wild type and BICD2^-/-^ RPE-1 cells. Although RPE-1 cells are non-transformed and p53^+/+^ cells, and we thus did not expect massive centriole number changes (Fava *et al*, 2017), we indeed observed small but significant differences in centriole numbers, with cells lacking BICD2 tending to have more centrioles than wild type cells. Thus, while only 3% of wild type cells showed more than the normal 4 centrioles in mitosis, 14% of each BICD2^-/-^ cell line had more than 4 centrioles, with some cells showing 6 or 7 centrioles, something that we never observed for wild type cells (Figure 6E, top left graph). This effect was more evident in transformed U2OS cells, see below.

As a more direct way to assess disengagement we used the mother centriole marker C-NAP1 (Fry *et al*, 1998). This protein accumulates at the proximal end of mother centrioles and is only detectable at daughter centrioles after they have disengaged from their partner. As a result in normal conditions early mitotic cells (i.e. with two non-disengaged diplosomes) have only two C-NAP1 foci, and more than two c-NAP1 foci indicate disengagement (Tsou & Stearns, 2006b; Tsou *et al*, 2009). We found significantly different numbers of C-NAP1-positive centrioles in BICD2^-/-^ cells as compared to their wild type counterparts, observing that while in mitosis only 11% of wild type RPE-1 cells showed more than two C-NAP1 foci, 48% and 34% of mitotic cells in the two BICD2^-/-^ cells lines showed more than two C-NAP1 foci, sometimes showing 6 or 7 of them (Figure 6E, top right graph). Being aware that BICD2^-/-^ cells have higher number of centrioles than wild type cells we also measured the ratio of C-NAP1 positive centrioles to the total number of centrioles to account for that difference. As expected most mitotic wild type cells had a 2:4 ratio, that is four centrioles two of them two positive for C-NAP1, with only 10% of them showing rations bigger than 2:4, i.e. more C-NAP1-positive centrioles per total number of centrioles. In contrast, we observed cells with ratios bigger than 2:4 in the 34% and 24% of cases in the two BICD2^-/-^ cell lines (Figure 6E, bottom graph).

We finally explored whether an abnormal BICD2 function could also affect centriole numbers in a transformed cell line. For this we downregulated BICD2 in U2OS osteosarcoma cells and quantified centriole numbers. We observed that 50% of cells transfected with a siRNA specific for BICD2 showed more than 4 centrioles in mitosis while only 20% of cells transfected with a control siRNA did, confirming that BICD2 downregulation can result in centriole amplification in different cell types (Figure 6F).

In order to assess the specificity of our observations and better understand them, we next expressed different GFP-tagged BICD2 polypeptides in BICD2^-/-^ cells and studied whether they could effectively rescue the observed phenotypes. As we did in Figure 6D we used STLC to arrest the cells in prometaphase. This had the advantage of resulting in a workable number of mitotic cells expressing the recombinant proteins. Figure 7A shows that the expression of full-length BICD2 (GFP-BICD2[1-820]) in both BICD2^-/-^ cell lines was able to rescue centriole separation in mitosis, resulting in intercentriolar distances similar to those observed in wild type cells. In contrast, neither GFP-BICD2[1-575], containing the N-terminal half of BICD2, nor GFP-BICD2[488-820] or GFP-BICD2[658-820], two C-terminal fragments of BICD2 that localized to the centrosomes (see Figure 4), were able to rescue the phenotype, essentially behaving as GFP alone. Thus, while BICD2 localizes to the centriole via its C-terminal region, at least part of its N-terminal domain, encompassing the CC1 and CC2 regions, is also essential for its role maintaining mother and daughter centrioles engaged in early mitosis. Studying the functional relevance of BICD2 N-terminus in similar experiments, we importantly observed that GFP-BICD2[65-820], lacking the CC1 box motif crucial to bind to dynein, and specifically its DLIC subunit (Lee *et al*, 2018), rescued the phenotype of BICD2^-/-^ cells as effectively as the wild type protein (Figure 7B). This suggested that the roles of BICD2 at the centriole were independent from its function as a dynein adaptor/activator, something that was confirmed by the observation that GFP-BICD2[272-820], a polypeptide lacking the whole N-terminal CC1 region and thus any dynein binding capability, behaved as the full-length protein, being able to rescue the disengagement phenotype. Note that both GFP-BICD2[65-820] and GFP-BICD2[272-820] are able to localize at centrosomes (see Figure 4) and are expressed at levels similar to those of wild type BICD2 (Supplementary Figure 8A). From these experiments we also concluded that the dynein-independent role of BICD2 at centriole involves the CC2, CC3 and CC4 regions of the protein.

**Figure 7.**
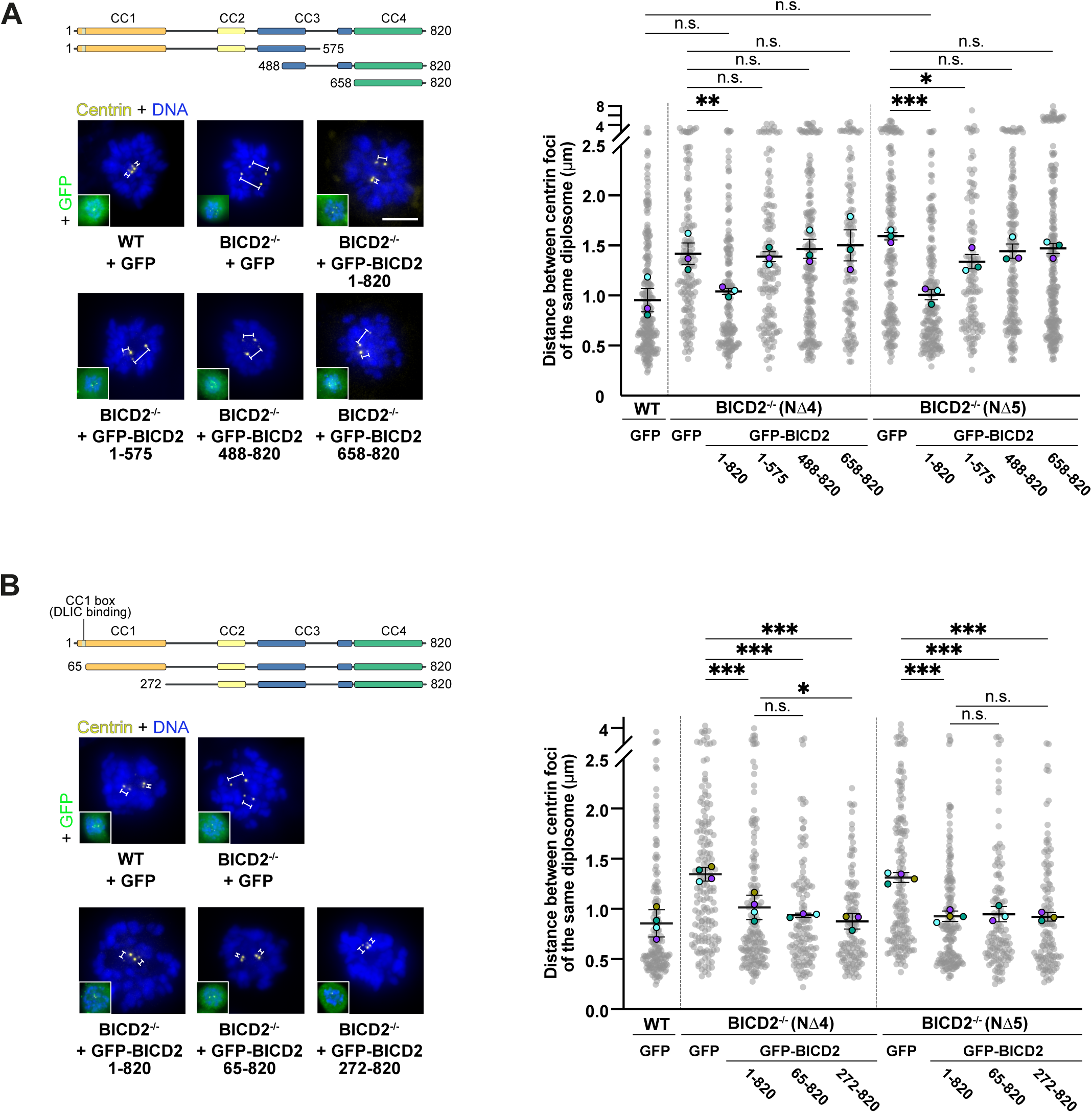
Recombinant full-length BICD2 but not BICD2 N- or C-terminal regions rescue abnormal intercentriolar distances resulting from the lack of endogenous protein. BICD2 forms that do not bind dynein are able to rescue intercentriolar distances similarly to the wild type protein. **A and B.** Distance between centrin foci of the same diplosome in wild type and BICD2^-/-^ RPE-1 cells expressing the indicated GFP-fusion proteins. Cells were transfected with the corresponding plasmids and subsequently incubated with STLC overnight before fixation (n=3 biological replicates, ∼50 diplosomes per replicate; individual replicate means plus mean ± SD of replicates are shown and statistical significance is analyzed using one-way ANOVA with post hoc analysis). Schematic representation of the different BICD2 forms and immunofluorescence images of example cells stained for centrin and DNA (A-D, F) or centrin, GFP and DNA are shown. Scale bar 5 µm. Expression of the different GFP-fusion proteins is shown in Supplementary Figure 8A.

We finally explored the mechanisms that could regulate the centriolar localization of BICD2 and result in a greatly diminished pool of protein at the centrosomes in mid/late mitosis, something that according to our data will result in mother-daughter disengagement and the licensing of centriole duplication. We have previously observed that most of BICD2 is phosphorylated in mitosis (Gallisà-Suñé *et al*, 2023). We thus wondered whether the modification of BICD2 could be interfering with its localization at the centriole. We noticed that the C-terminal region of BICD2 is phosphorylated *in vivo* at several sites, and that two of the most frequently identified phosphoresidues in high throughput proteomic studies, Thr821 and Ser823 in human BICD2, are located at the extreme C-terminus of the protein (see PhosphositePLus (https://www.phosphosite.org; the homologous residues in mice, Ser817 and S819, have also been shown to be phosphorylated *in vivo*). Importantly, Thr821 can be phosphorylated by the main mitotic regulator CDK1 (Fagiewicz *et al*, 2022) and the phosphorylation levels of both Thr821 and Ser823 are diminished upon PLK1 inhibition (Kettenbach *et al*, 2011), suggesting a role for these kinases in the modification of the residues in mitosis. We thus tested whether their modification interfered with the ability of the C-terminal region of the protein to localize to the centrosome. As can be seen in Figure 8A, the presence of phosphomimetic residues at the extreme C-terminus of BICD2 (S817D, S819D in the mouse protein used in this study) significatively affected the ability of both a C-terminal fragment of BICD2 (GFP-BICD2 [658-820]) and the full-length protein (GFP-BICD2 [1-820]) to localize at the centrosome, both in wild type and BICD2^-/-^ RPE-1 cells. This supported the hypothesis that the phosphorylation of BICD2 C-terminus is able to control BICD2 centriolar localization. Moreover, in contrast to wild type BICD2, the phosphomimetic form of the protein (BICD2 S817D, S819D) failed to rescue the abnormal centriole distances observed in BICD2^-/-^ cells (Figure 8B).

**Figure 8.**
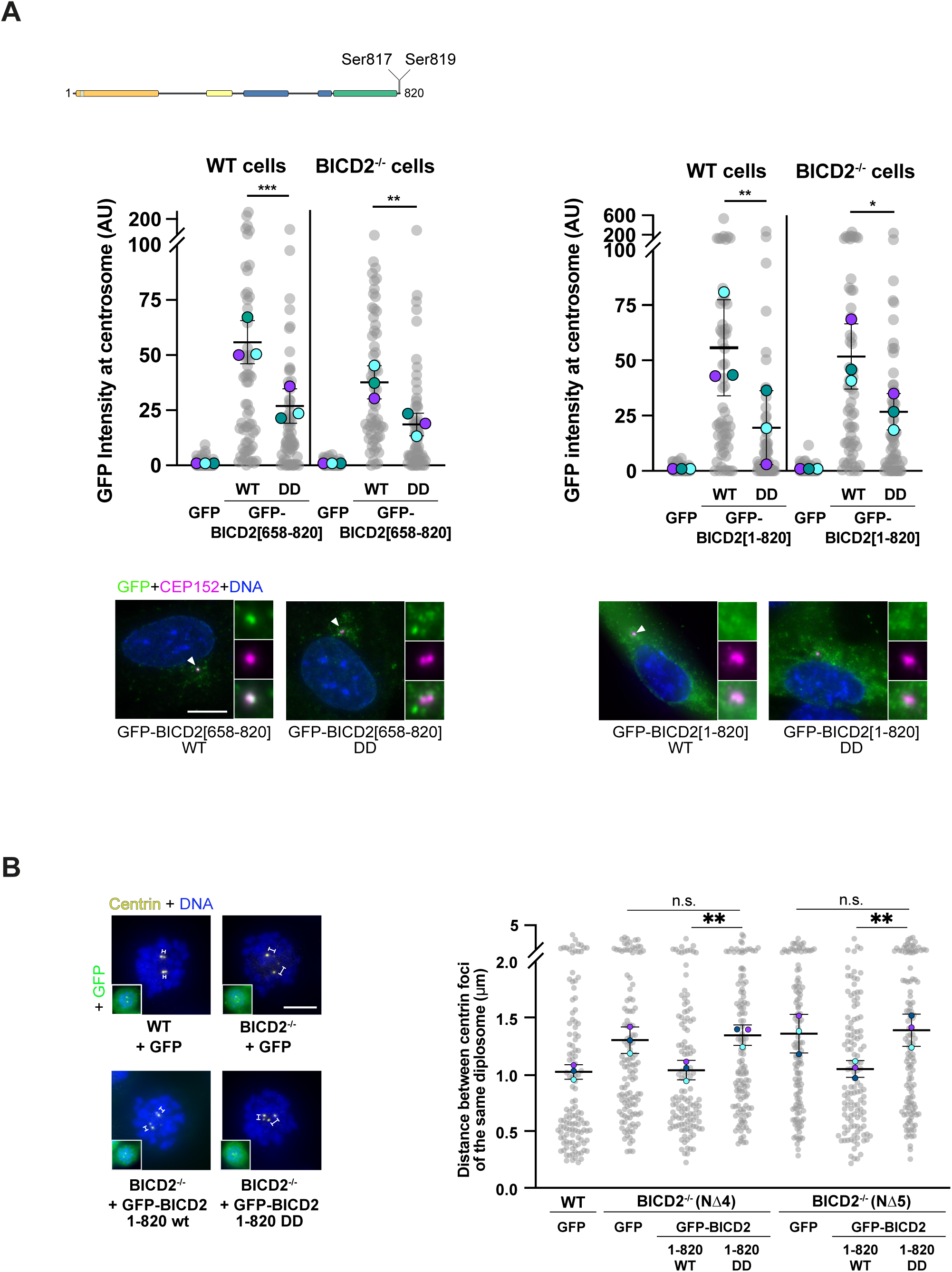
Recombinant phosphomimetic forms of BICD2 fail to localize to the centriole and to rescue abnormal intercentriolar distances resulting from the lack of endogenous protein. **A.** *Top*, schematic representation of BICD2 showing the location of Ser817 and Ser819. *Middle*, centrosomal intensities of the GFP signal in both wild type and BICD2^-/-^ RPE-1 cells expressing the indicated recombinant proteins, fixed and stained for GFP, CEP152 as a centrosomal marker, and DNA. G1/S cells transfected with low levels of GFP constructs were chosen (n=3 biological replicates, 20 cells per replicate; individual replicate means plus mean ± SD of replicates are shown; statistical significance is analyzed using one-way ANOVA with post hoc analysis). Example immunofluorescence images of BICD2^-/-^ cells are shown below the graphs. Scale bar 10 µm. **B.** Distance between centrin foci of the same diplosome in wild type and BICD2^-/-^ RPE-1 cells expressing the indicated GFP-fusion proteins. Cells were transfected with the corresponding plasmids and subsequently incubated with STLC overnight before fixation (n=3 biological replicates, 40 diplosomes per replicate; individual replicate means plus mean ± SD of replicates are shown; statistical significance is analyzed using one-way ANOVA with post hoc analysis). Example immunofluorescence images are shown. Scale bar 5 µm. *BICD2 DD*, BICD2 S817D, S819D. Expression of the different GFP-fusion proteins is shown in Supplementary Figure 8C.

## Discussion

Our results reveal a previously unknown function for BICD2, a protein best known as one of the adaptors/activators of the microtubule motor dynein (Hoogenraad *et al*, 2001; Hoogenraad & Akhmanova, 2016). BICD2 is a coiled-coil rich dimer (see Figure 4A) whose functions are multiple and cell-cycle dependent. In interphase it acts as a classical dynein adaptor, linking the motor to Golgi and exocytotic vesicles that it transports though its interaction with the small G-protein Rab6 (Matanis *et al*, 2002; Grigoriev *et al*, 2007). In G2 BICD2 is phosphorylated and switches cargos, binding to the nucleoporin RanBP2/NUP358 and to the outer nuclear membrane protein Nesprin-2. This results in the recruitment of active dynein to the nuclear envelope and is instrumental for the correct separation of centrosomes in G2 and early M (Splinter *et al*, 2010; Gonçalves *et al*, 2020; Gallisà-Suñé *et al*, 2023). We now show that additionally to those functions, BICD2 has a dynein-independent role at the centrosome, where it localizes around the mother and close to the daughter centriole and contributes to maintain mother-daughter engagement until late mitosis, when it loses its centriolar localization.

A centrosomal localization for BICD2 is suggested by published immunofluorescence microscopy data from different authors (Hoogenraad *et al*, 2001; Quarantotti *et al*, 2019) and supported by proteomic studies that indicate that it may interact with a number of centrosomal proteins (e.g. (Splinter *et al*, 2010; Gupta *et al*, 2015; Redwine *et al*, 2017; Carden *et al*, 2023)). We confirm this localization using different antibodies, techniques and cell lines. BICD2 accumulates at a proximal region of the mother centriole coinciding with the site of daughter centriole formation and forms a usually discontinuous ring around it. It begins to accumulate at this region during the G1 phase of the cell cycle, coinciding with centriole to centrosome conversion, and its localization partially overlaps and temporarily follows that of CEP152. The centriolar localization of BICD2 results from the ability of its C-terminal region to accumulate at the organelle, and accordingly recombinant polypeptides comprising only the last coiled-coil region of BICD2 (CC4) are strikingly centrosomal when expressed in mammalian cells. Shorter C-terminal regions partially lose this localization, something that might indicate that a dimeric form is needed for localization, as possibly these forms may not form dimers so efficiently. Alternatively, the centriolar localization motif in BICD2 may be internal to CC4 and not present in short C-terminal polypeptides. Further work should clarify this and identify the centriolar proteins that interact with BICD2. At mature centrioles, the components of the proximal torus formed by CEP57, CEP63 and CEP152 make good candidates. Our superresolution images indicate that a pool of BICD2 has a localization that partially coincides with that of SAS-6, suggesting that BICD2 might also interact with daughter centriole components, making cartwheel proteins obvious candidates to look at. In fact, SAS-6 has been reported as a BICD2 interactor by proximity-label MS (Redwine *et al*, 2017). Further investigations will also clarify the importance of the unstructured loop that is predicted to protrude from the CC4 region of BICD2 and explain how it may promote the organization of higher order annular structures, e.g. by multimerizing through non-specific interactions and facilitating the interaction between BICD2 dimers.

Our data shows that BICD2 is key for the maintenance of centriole engagement in the G2 and M phases of the cell cycle. Mother and daughter centrioles do not depend on BICD2 to stay associated in S phase, suggesting that different engagement mechanisms exist during the progression through the centriolar cycle, maybe related to the fact that at this phase the daughter (pro)centriole is still growing and firmly attached to the mother through the cartwheel. Importantly, we show that BICD2 forms lacking the ability to interact with dynein exhibit localization patterns and functional properties similar to their wild-type counterparts. Thus, BICD2 keeps mother and daughter centrioles engaged independently of its ability to bind the dynein motor complex. How does it perform this function? BICD2 is a dimer (probably a cotranslational one (Bertolini *et al*, 2021)), and in its dimeric form has a rod-like extended structure with a length predicted to be of minimally ∼50 nm, and more than 100 nm if completely extended (see Figure 4A; see also (Fagiewicz *et al*, 2022; Gallisà-Suñé *et al*, 2023) for electron microscopy images of BICD2 dimers). We thus hypothesize that taking advantage of this geometry it might physically bridge mother and daughter centrioles by interacting with specific components of both of these structures through sequences in the CC2/CC3 and CC4 regions of the protein.

As expected from a protein controlling engagement and ultimately licensing, BICD2 amounts at the centriole are dynamic and markedly diminish in late G2 and M, with only a small pool of protein remaining close to the mother-daughter centriole interface until late mitosis, when disengagement occurs. Considering our findings with phosphomimetic BICD2 mutants, and given that most BICD2 is phosphorylated in mitosis (Gallisà-Suñé *et al*, 2023), we propose that phosphorylation of the extreme C-terminus of BICD2 at Thr821 and Ser823 by mitotic kinases such as CDK1 and/or PLK1, both previously shown to be able to modify BICD2 (Fagiewicz *et al*, 2022; Gallisà-Suñé *et al*, 2023), may disrupt the binding of the protein to its receptors during late G2/M, especially those localized at the mother centriole. In this manner phosphorylation could contribute to centriole disengagement by removing most of BICD2 from the centrioles. And dephosphorylation during mitotic exit or early G1 might result in the (re)accumulation of BICD2 at mature centrioles during centriole to centrosome conversion. Phosphorylation alone may suffice to remove enough BICD2 to result in disengagement in late mitosis. Alternatively, the last remnants of centriolar BICD2, that we observe close to the daughter centriole, might be removed by separase cleavage. This will be the subject of future studies, but we would like to note that BICD2 contains in its CC3 domain a putative separase cleavage motif of the form TXEXXRX (Hauf *et al*, 2001).

Finally, it is important to note that BICD2^-/-^ RPE-1 cells show normal levels of pericentrin in mitosis. This indicates that, at least in RPE-1 cells, pericentrin removal and PCM disassembly are not requisites for centriole disengagement. Although it has been shown that pericentrin cleavage can result in disengagement and that expression of non-cleavable forms of the protein suppresses normal disengagement, our results do not agree with the suggestion that separase-dependent cleavage of pericentrin is necessary for this process to occur during mitosis (Matsuo *et al*, 2012; Lee & Rhee, 2012; Kim *et al*, 2019). In fact, BICD2^-/-^ cells show signs of centriole disengagement in G2 or early M, when pericentrin could not be cleaved, being separase inactive. Pericentrin may have a complementary role to that of BICD2 keeping mother and daughter centrioles engaged, and its downregulation from the PCM during mitotic exit may facilitate disengagement induced by the loss of BICD2. However, we show that the removal of the protein is in itself not necessary for disengagement to occur. We would like to additionally note that the downregulation of pericentrin either does not induce centriole disengagement or only minimally affects it (Lee & Rhee, 2012; Kim *et al*, 2019), and only complete deletion of the protein has been shown to result in premature centriole separation in mitosis (Kim *et al*, 2019). The removal of pericentrin is expected to have a massive effect on PCM structure, and the separation of the centrioles observed by previous authors in response to it may thus be the result of creating abnormal PCM-free diplosomes and reflect a condition that is not physiologically relevant in normally cycling cells, which always retain some level of PCM (including pericentrin) surrounding their mother centrioles.

In conclusion, we have proved that a pool of BICD2 resides at the centrosome and that its localization there is disrupted in late mitosis, probably in response to phosphorylation. Centrosomal BICD2 is necessary to maintain mother and daughter centrioles together until the end of mitosis, and its removal from the organelle results in centriole disengagement, and thus, centriole duplication licensing. Therefore, BICD2 has a main role in centriole engagement during the centrosome cycle.

## Experimental Procedures

### Mammalian expression constructs and siRNAs

pEGFP-C2-BICD2, containing the cDNA for mouse BICD2, was a gift from Anna Akhmanova (Utrecht University) and is described in (Hoogenraad *et al*, 2001). Fragments and mutant forms of BICD2 were produced directly by PCR or using QuickChange Lighting Site-Directed Mutagenesis kit (Agilent Technologies) according to the manufacturer’s instructions using the following primers and the appropriate reverse complements. All clones were sequenced and expression of polypeptides assessed by western blot.

BICD2 [1-575]:

5’-GAGGGCCGCGGGTGACGGTCACCTGTC-3’;

BICD2 [576-820]:

5’-GATCTCGAGAGAATTCGCCACCCGCCGGTCACCTGTCCTCTTG-3’;

BICD2 [65-820]:

5’-GAGAGAATTCGCCACCGAGGTGGACTATGAGG-3’

BICD2 [272-820]:

5’-AAAAGGATCCTATACCAGCCACCTGCAGGTC-3’

5’-AAAAGGATCCCTACAGGCTCGGTGAGGCTGGCTTG-3’

BICD2 [488-820]:

5’-GATCTCGAGAGAATTCGCCACCTCTCTGCTGGAGAAGGCTAGCC-3’

BICD2[488-820, Δ547-620]:

5’-CACCATGTGTGCATGTGCATGAACATCTACAACCTG-3’

BICD2 [576-820[:

5’-GATCTCGAGAGAATTCGCCACCCGCCGGTCACCTGTCCTCTTG-3’

BICD2 [620-820]: 5’-GAATTCGCCACCATGCCTATGAACATCTAC-3’

BICD2 [658-820]: 5’-GAATTCGCCACCATGGGCCCTGCTGTGGAC-3’

BICD2 [754-820]:

5’-CTCAGATCTCGAGAGAATTCATCACACAGCTGGATGAGATG-3’

BICD2[802-820]:

5’-AATTCACCCGCAGGGGCCGCAGCAAGGCCGCCTCCAAGGCCANGCCAGCCTCACCGAGCCTGTAGG-3’

BICD2 [488-798]: 5’-CTGGAGCTGGACTAAGAACAGACCCGC-3’

BICD2 [576-798]: 5’-CTGGAGCTGGACTAAGAACAGACCCGC-3’

BICD2 [488-575]: 5’-GAGGGCCGCGGGTGACGGTCACCTGTC-3’

BICD2 S817D, S819D: 5’-CCAAGCCAGCCGATCCGGACCTGTAGAGTAG-3’

In order to engineer BICD2^-/-^ cells using CRISR/Cas9 genome editing, we cloned the following oligonucleotide into pSpCas9(BB)-2A-GFP (PX458, from Feng Zhang; Addgene plasmid #48138 (Ran *et al*, 2013)): 5’-AAACCCAAGGAGCAGTACTACGTGC-3’, containing a gRNA corresponding to a sequence in exon 2 of the human BICD2 gene.

For endogenous BICD2 knockdown we used the following siRNA sequence: 5’-GGAGCUGUCACACUACAUGUU-3’. A siRNA against luciferase with the following sequence was used as control: 5’-CGUACGCGGAAUACUUCGA-3’.

### Cell culture and transfection

hTERT RPE-1 cells (ATCC CRL-4000) and U2OS cells (ATCC HTB-96) were grown in DMEM/F12 or DMEM media (both Thermo Fisher), respectively, supplemented with 10% foetal bovine serum (Thermo Fisher), 2mM L-glutamine (Thermo Fisher) and 100 U/mL penicillin/ 100 ug/mL streptomycin (Thermo Fisher).

RPE-1 cells were transfected using TransfeX (ATCC) and Opti-MEM media (Thermo Fisher) according to manufacturer’s instructions. For RNAi, U2OS cells were transfected with siRNAs using RNAiMAX (Thermo Fisher) and after 72 h re-transfected. 48 h after that cells were fixed with cold methanol for immunofluorescence or harvested for western blot.

For rescue experiments, wild type and BICD2^-/-^ RPE-1 cells were transfected with the indicated plasmids. After 7 h, 5 µM of STLC ((+)-S-Trityl-L-cysteine, Sigma) was added to the media. Cells were incubated for 16 h and fixed with cold methanol for immunofluorescence.

### Generation of CRISPR knockout cell lines

BICD2^-/-^ RPE-1 cells were transfected using Lipofectamine 2000 (Thermo Fisher) according to manufacturer’s instructions with the corresponding pSpCas9(BB)-2A-GFP plasmid and, 48 hours following transfection, cells positive for GFP were sorted in a SORTER FUSION II and seeded in 96-well plates. Single cell clones were expanded during 2-4 weeks and screened by western blot for BICD2 protein levels. Relevant clones were subsequently sequenced.

### Antibodies

Primary antibodies used were as follows:

anti-acetylated tubulin: mouse IgG2b 6-11B (Sigma/Santa Cruz)

anti-BICD2: mouse IgG2b GT10811/GTX631994 (GeneTex); rabbit polyclonal ab117818 (Abcam); rabbit polyclonal #2293 (Hoogenraad *et al*, 2001); rabbit polyclonal NBP1-81488 (Novus Biologicals); rabbit polyclonal A10292 (Abclonal)

anti-centrin: mouse IgG2A 20H5 (Merck)

anti-centrobin: mouse IgG1 (Ogungbenro *et al*, 2018)

anti-CEP152: mouse IgG1 GT1315/GTX630984 (GeneTex)

anti-CEP164: mouse IgG2A sc-515403 (Santa Cruz)

anti-C-NAP1: rabbit polyclonal 14498-1-AP (Proteintech)

anti-GFP: rabbit polyclonal A6455 (Invitrogen); mouse IgG2a 3E6 (Lifetech/ThermoFisher)

anti-GM130: mouse IgG1 610822 (BD Transduction Laboratories)

anti-PCM-1: mouse IgG1 sc-398365 (Santa Cruz)

anti-pericentrin: rabbit polyclonal 4448 (AbCam)

anti-SAS-6: mouse IgG2b sc-81431 (Santa Cruz)

Secondary antibodies used were as follows (Alexa Fluor conjugates, Thermo Fisher; HRP conjugates, R&D):

Alexa Fluor 647 goat anti-rabbit IgG (H+L) A-21244

Alexa Fluor 488 goat anti-rabbit IgG (H+L) A-11008

Alexa Fluor 647 goat anti-mouse IgG1 A-21240

Alexa Fluor 488 goat anti-mouse IgG1 A-21121

Alexa Fluor 568 goat anti-mouse IgG1 A-21124

Alexa Fluor 488 goat anti-mouse IgG2a A-21131

Alexa Fluor 647 goat anti-mouse IgG2a A-21241

Alexa Fluor 568 goat anti-mouse IgG2b A-21144

Alexa Fluor 647 goat anti-mouse IgG2b A-21242

Goat anti-mouse IgG-HRP HAF007

Goat anti-rabbit IgG-HRP HAF008

### Immunofluorescence microcopy

Cells were grown on poly-L-Lysine coated coverslips to subconfluence. After the indicated treatments, cells were rinsed in cold PBS and fixed with methanol at −20 °C for at least 1h. After three washes in PBS, cells were incubated for 20min in blocking buffer (3% BSA, 0.1% Triton X-100, 0.02% NaN_3_ in PBS) and stained using the indicated primary antibodies diluted in blocking buffer for 1 h at room temperature. After two washes with PBS, cells were incubated with the secondary antibodies diluted in blocking buffer for 20-40 min at room temperature. Slides were mounted in ProLong Glass Antifade Mountant with NucBlue mounting media (Thermo Fisher) to stain DNA and stored in a dark chamber overnight.

Three and four channel Z-stack images were acquired with an ORCA-Flash 4.0 LT+ Digital CMOs camera (Hamamatsu C11440) in a Leica DMi8 inverted microscope equipped with 63×/1.40NA oil immersion lens and the standard LAS software. Maximum projections of z-stacks were produced using FIJI software (ImageJ).

Three-dimensional structured illumination microscopy (3D-SIM) data was acquired on a LSM 880 Zeiss confocal microscope equipped with an Elyra PS1 module and a piezo stage. Image acquisition was done with a 100x/1.46NA oil Dic M27 lens. Grid rotations were restricted to three to avoid bleaching. SIM images were reconstructed with the ZEN black software from Zeiss. The fluorescent dyes were excited using 488 nm (30% of 200 mW laser source), 561 nm (20% of 200 mW laser source), and 642 nm (7% of 500 mW laser source) lasers. DNA staining was excited using a 405nm (50% of 50 mW laser source). The emitted light was collected through 420-480nm (405nm excitation) 495–575 nm (488 nm excitation), 570–650 nm (561 nm excitation), and 655 nm-above (642 excitation) emission filters. When indicated, maximum projections or orthogonal views of the z-stack were produced using FIJI software (ImageJ).

### Quantification and statistical analysis

Measurements of intensities and distances were done using FIJI software (ImageJ). Graphics and statistical analysis were done using Prism 9 software. Details of the number of measurements and biological replicates, as well as of the statistical tests used in each case are given in the corresponding figure legends (n.s. not significant, P > 0.05; * P < 0.05; ** P < 0.01; *** P < 0.001).

## Abbreviations

PCM: pericentriolar material.
3D-SIM: three-dimensional structured illumination microscopy.

## Acknowledgements

We are grateful to Anna Akhmanova (Utrecht University) for BICD2 plasmids and BICD2 (#2293) antibodies and to Ciaran Morrison (University of Galway) for centrobin antibodies. Work was funded by the Ministerio de Ciencia e Innovación (MICINN) and the Agencia Estatal de Investigación (AEI) from Spain (MCIN/ AEI /10.13039/ 501100011033/ FEDER “Una manera de hacer Europa”) through Plan Nacional de I+D grant PID2021-127045NB-I00 (to J.R.) and by network grant SGR-Cat-2021 00421 Dynamic Architecture of Life, LifeArchCAT (AGAUR, Generalitat de Catalunya). N.G.-S. was a recipient of FPI Fellowship BES-2015-072446 from MICINN.

## Notes

### Competing Interest Statement

The authors have declared no competing interest.

### Summary of Updates

Minor changes. Cosmetic improvements in the figures and supplemental figures.

