## Supplementary Figures for "BICD2 is a centriolar protein that controls mother-daughter centriole engagement and the licensing of duplication"

##### Supplementary Figure 1 (related to Figure 1).

Immunofluorescence images of RPE-1 cells in different phases of the cell cycle stained for BICD2 (GT10811 antibody), the centrosome marker CEP152, the Golgi marker GM130 and DNA. Scale bar 10 µm.

##### Supplementary Figure 2 (related to Figure 1).

Immunofluorescence images of RPE-1 (A) and U2OS (B) cells stained with different antibodies against BICD2, CEP152 plus DNA. Bottom graph, quantification of the centrosomal intensity of the BICD2 signal in U2OS cells in different phases of the cell cycle. Cells were synchronized in G1 by a double thymidine block, released into the cell cycle and images were acquired at different times corresponding to the indicated phases (20 centrosomes per phase; means ± SD are shown). Scale bar 10 µm.

##### Supplementary Figure 3 (related to Figure 1).

**A.** Control RPE-1 cells and cells incubated on ice for 30 minutes were fixed and stained for BICD2 (GT10811 antibody), centrin and CEP152 as centrosome markers, plus DNA. Centrosomal BICD2 was quantified in cells with two centrosomes (n=3 biological replicates, 20 centrosomes per replicate; values normalized to the mean values of control cells; individual replicate means plus mean ± SD of replicates are shown; statistical significance analyzed using an unpaired t-test with a two-tailed P value). Bottom panels show cells stained for BICD2 (#2293 antibody), centrin plus acetylated tubulin, confirming cold-induced microtubule depolymerization. Scale bar 10 µm.

**B.** Immunofluorescence images of G1/S and mitotic RPE-1 cells stained with antibodies against BICD2 (GT10811), pericentrin (PCNT) and PCM-1 plus DNA.

#### Supplementary Figure 4 (related to Figure 2)

**A.** 3D-SIM images of centrosomes of RPE-1 cells stained using two different BICD2 antibodies (GT10811 and ab117818), plus CEP152 antibodies. Maximum projections of z-stacks are shown. Scale bar 0.5 µm.

**B.** As in A, cells stained for BICD2 (GT10811 antibody), the appendage protein and mother (M) centriole marker CEP164 and the daughter (D) centriole marker centrobin.

**C.** As in A, cells stained for BICD2 (GT10811 antibody), CEP164 and C-NAP1. Maximum projections of z-stacks are shown in the left panels; single z-planes plus an orthogonal view of the z-stack are shown in the left.

**D.** 3D-SIM images of centrosomes of cells in G1/S, G2 and mitosis stained for BICD2 (GT10811 antibody), pericentrin and DNA.

#### Supplementary Figure 5 (related to Figure 2)

3D-SIM images of centrosomes of RPE-1 cells in the indicated phases of the cell cycle stained for BICD2 (GT10811 antibody), CEP152 and centrin. Maximum projections of z-stacks are shown. An overview image of the cell including DNA staining is additionally shown for mitotic cells. Scale bar 0.5 µm.

#### Supplementary Figure 6 (related to Figure 2)

**A.** 3D-SIM images of centrosomes of U2OS cells stained for BICD2 (ab117818 antibody); CEP152 identifies the mother centriole (M) while SAS-6 labels the daughter centriole (D). Maximum projections of z-stacks are shown. Scale bar 0.5 µm.

**B.** As in A, showing examples of cells in the indicated phases of the cell cycle. An overview image of the cell including DNA staining is additionally shown for mitotic cells. Note that in G1 only one centriole (the CEP152-positive mother) is visible, as the daughter has lost SAS6 during mitosis.

#### Supplementary Figure 7 (related to Figure 2).

Image scanning microscopy (Airyscan) images of centrosomes of U2OS cells stained for BICD2 (#2293 antibody), CEP164 and centrobin. CEP164 identifies the mother (M) centriole while centrobin acts as a daughter (D) centriole marker. Note that in some cases the younger mother centriole retains centrobin. Maximum projections of z-stacks are shown. An overview image of the cell including DNA staining is additionally shown. Scale bar 0.5 µm.

#### Supplementary Figure 8 (related to Figures 4, 6, 7 and 8).

A. Expression of the indicated GFP-fusion proteins in RPE-1 cells after transfection, as determined by Western blot (*W*) of total cell extracts.

B. BICD2 levels in U2OS transfected with control (*C*) and BICD2 siRNAs as determined by western blot of total cell extracts with anti-BICD2 antibodies (ab117818).

C. Expression of the indicated GFP-fusion proteins in RPE-1 cells after transfection, as determined by Western blot (*W*) of total cell extracts. *DD*, S817D, S819D.

In all cases Coomassie staining of the membrane is shown to assess protein loading.

### SUP. FIGURE 1

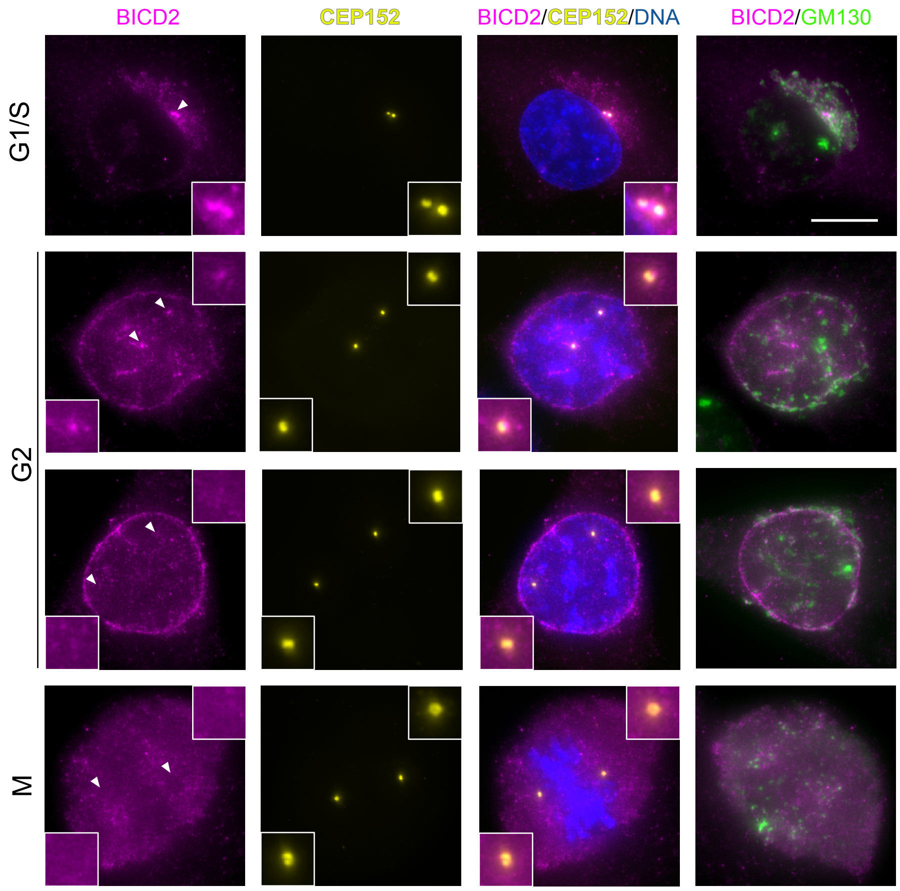

### SUP. FIGURE 2

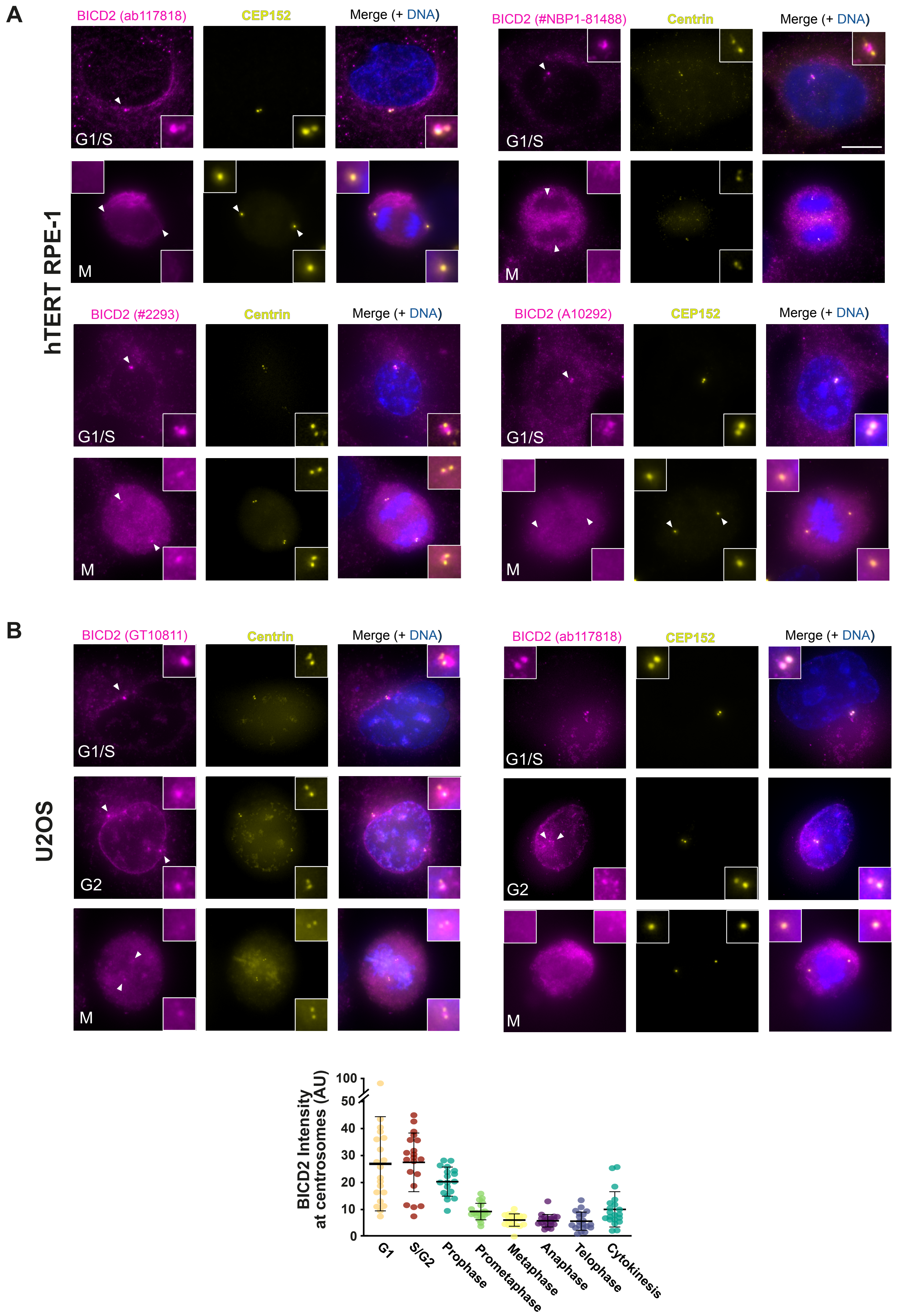
A
hTERT RPE-1
BICD2 (ab117818)
CEP152
Merge (+ DNA)
BICD2 (#NBP1-81488)
Centrin
Merge (+ DNA)
G1/S
M
BICD2 (#2293)
Centrin
Merge (+ DNA)
BICD2 (A10292)
CEP152
Merge (+ DNA)
B
U2OS
BICD2 (GT10811)
Centrin
Merge (+ DNA)
BICD2 (ab117818)
CEP152
Merge (+ DNA)
G1/S
G2
M
BICD2 Intensity at centrosomes (AU)
100
50
40
30
20
10
0
G1
S/G2
Prophase
Prometaphase
Metaphase
Anaphase
Telophase
Cytokinesis

### SUP. FIGURE 3

A

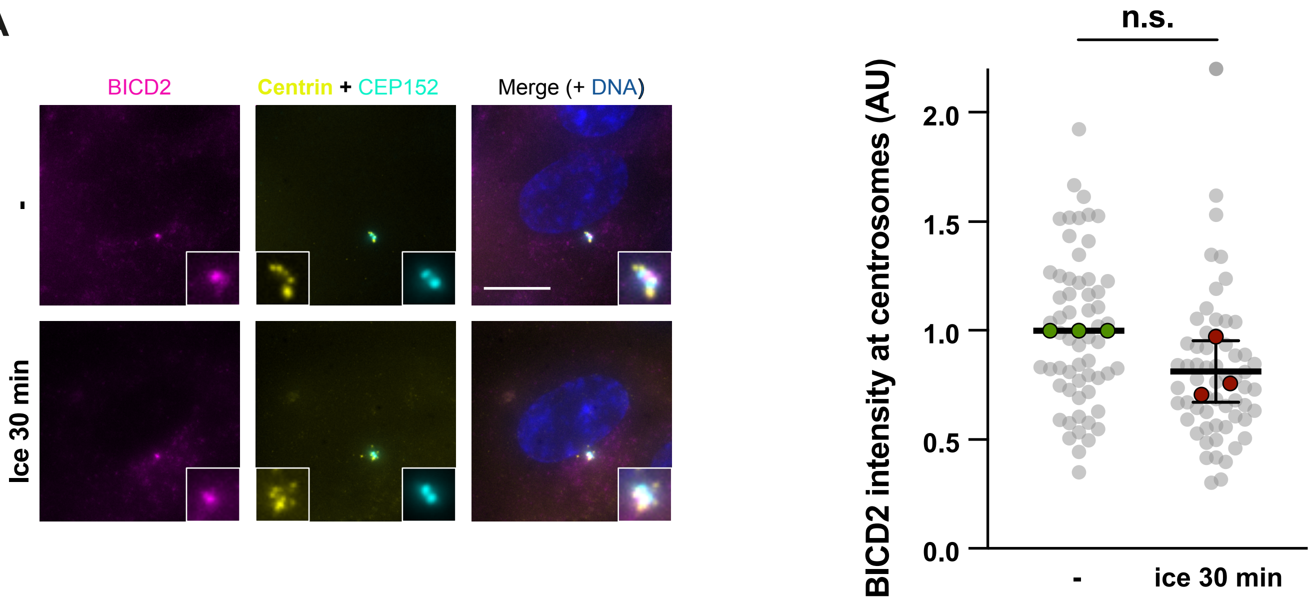


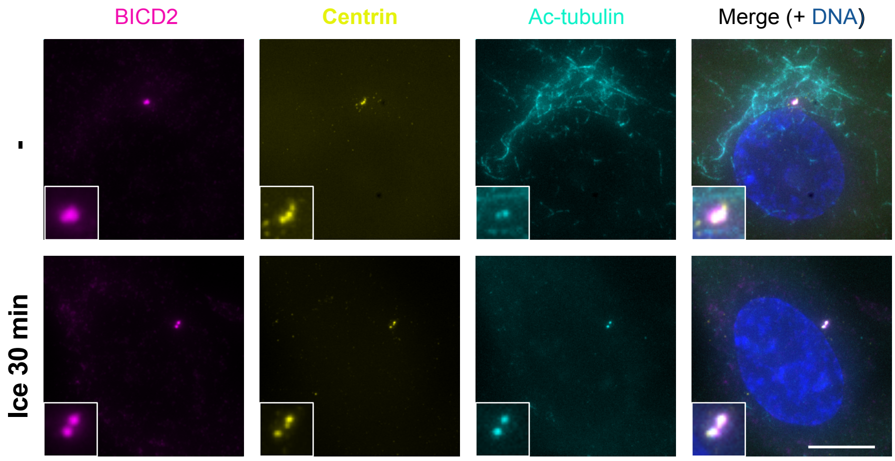


B

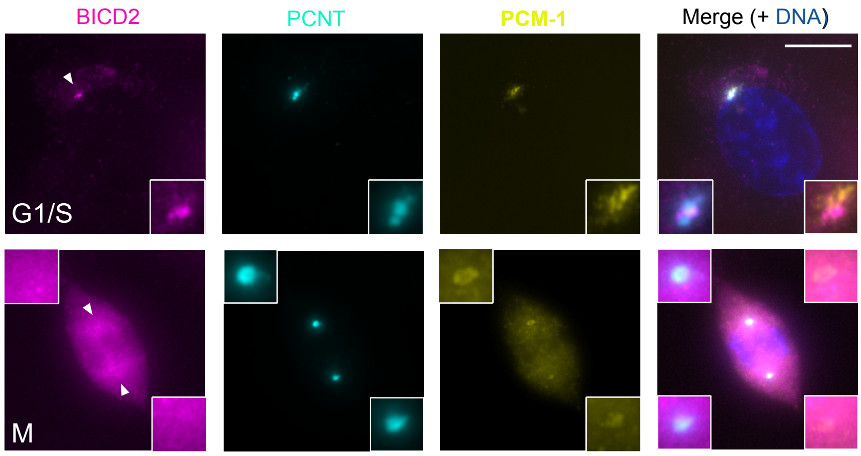

### SUP. FIGURE 4

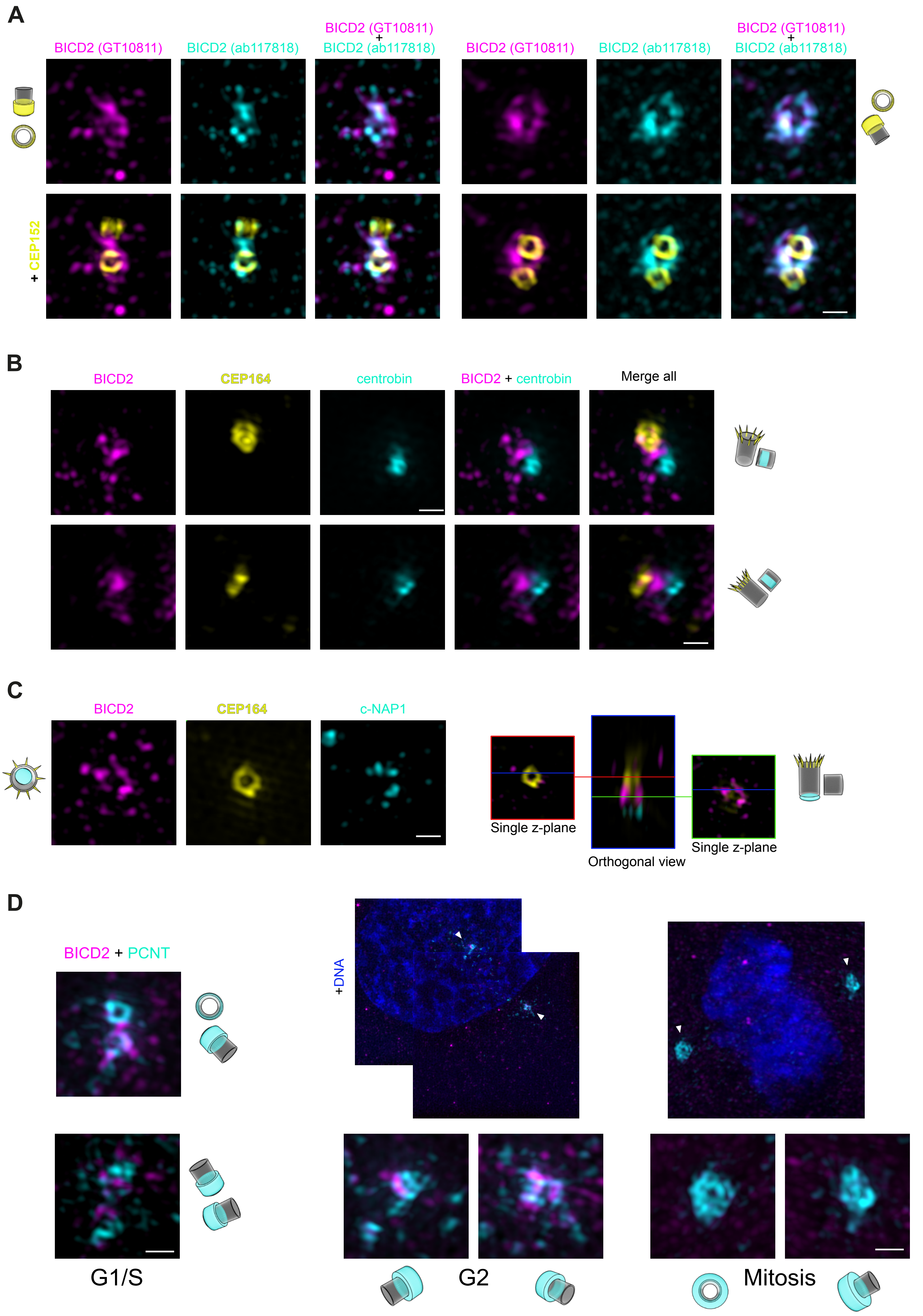
A
BICD2 (GT10811)
BICD2 (ab117818)
BICD2 (GT10811) + BICD2 (ab117818)
+ CEP152
B
BICD2
CEP164
centrobin
BICD2 + centrobin
Merge all
C
BICD2
CEP164
c-NAP1
Single z-plane
Orthogonal view
Single z-plane
D
BICD2 + PCNT
+DNA
G1/S
G2
Mitosis

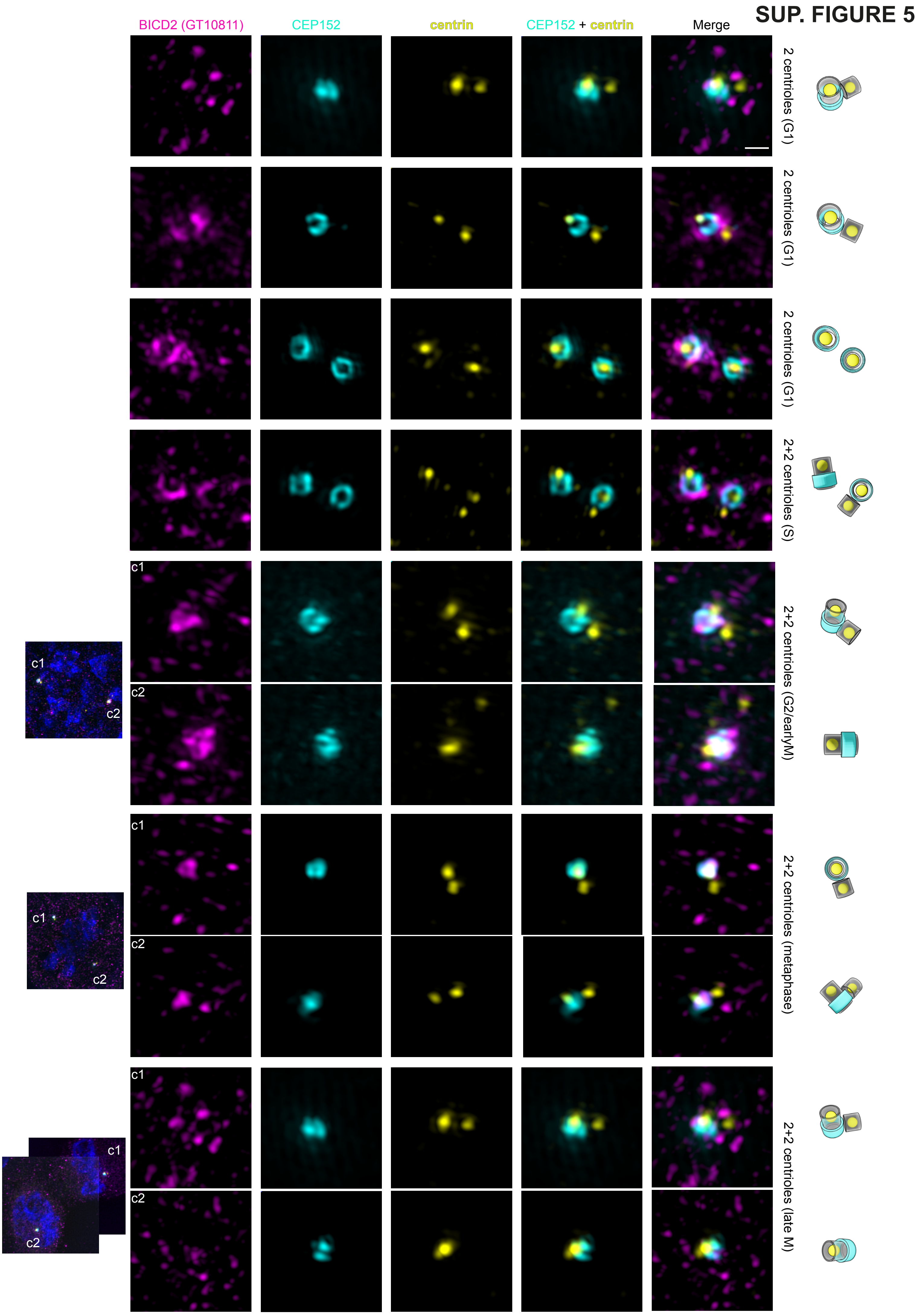
SUP. FIGURE 5
BICD2 (GT10811)
CEP152
centrin
CEP152 + centrin
Merge
2 centrioles (G1)
2 centrioles (G1)
2 centrioles (G1)
2+2 centrioles (S)
c1
c2
2+2 centrioles (G2/earlyM)
c1
c2
2+2 centrioles (metaphase)
c1
c2
2+2 centrioles (late M)

### SUP. FIGURE 6

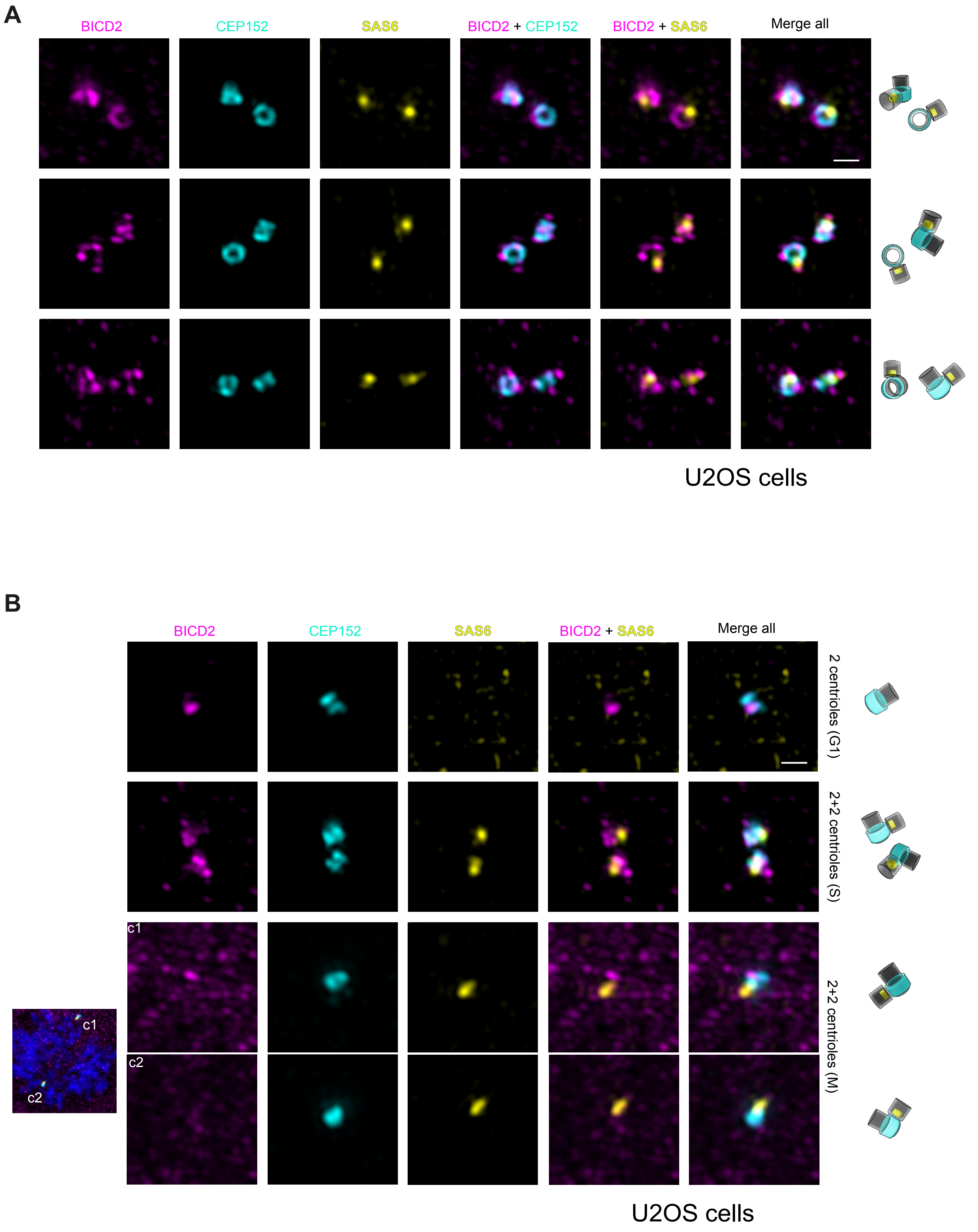
A
BICD2
CEP152
SAS6
BICD2 + CEP152
BICD2 + SAS6
Merge all
U2OS cells
B
BICD2
CEP152
SAS6
BICD2 + SAS6
Merge all
2 centrioles (G1)
2+2 centrioles (S)
2+2 centrioles (M)
c1
c2
U2OS cells

#### SUP. FIGURE 7

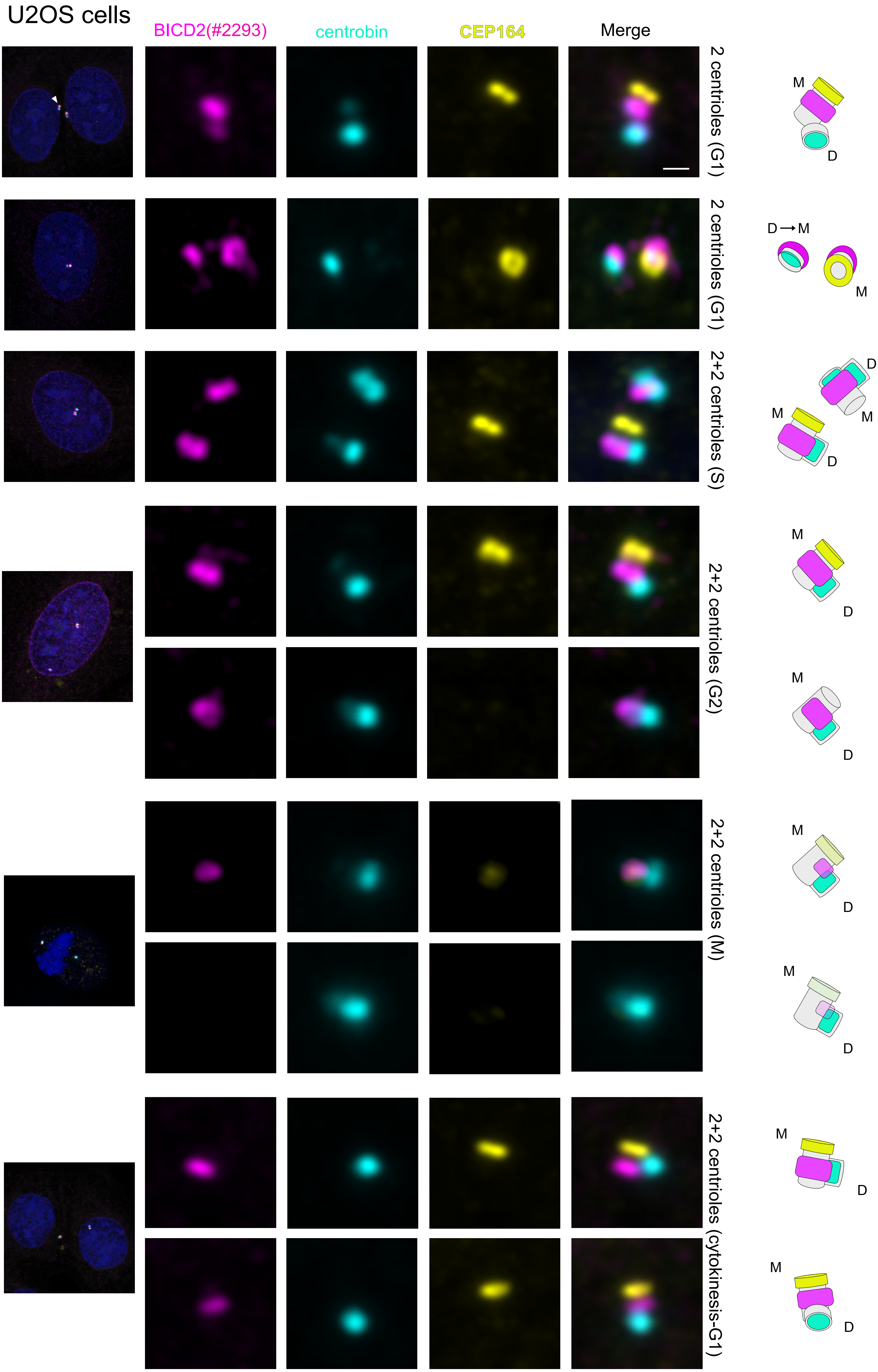
U2OS cells
BICD2(#2293)
centrobin
CEP164
Merge
2 centrioles (G1)
2 centrioles (G1)
2+2 centrioles (S)
2+2 centrioles (G2)
2+2 centrioles (M)
2+2 centrioles (cytokinesis-G1)
M
D
D→M
M
D
M
M
D
M
D
M
D
M
D
M
D
M
D
M
D

**SUP. FIGURE 8**

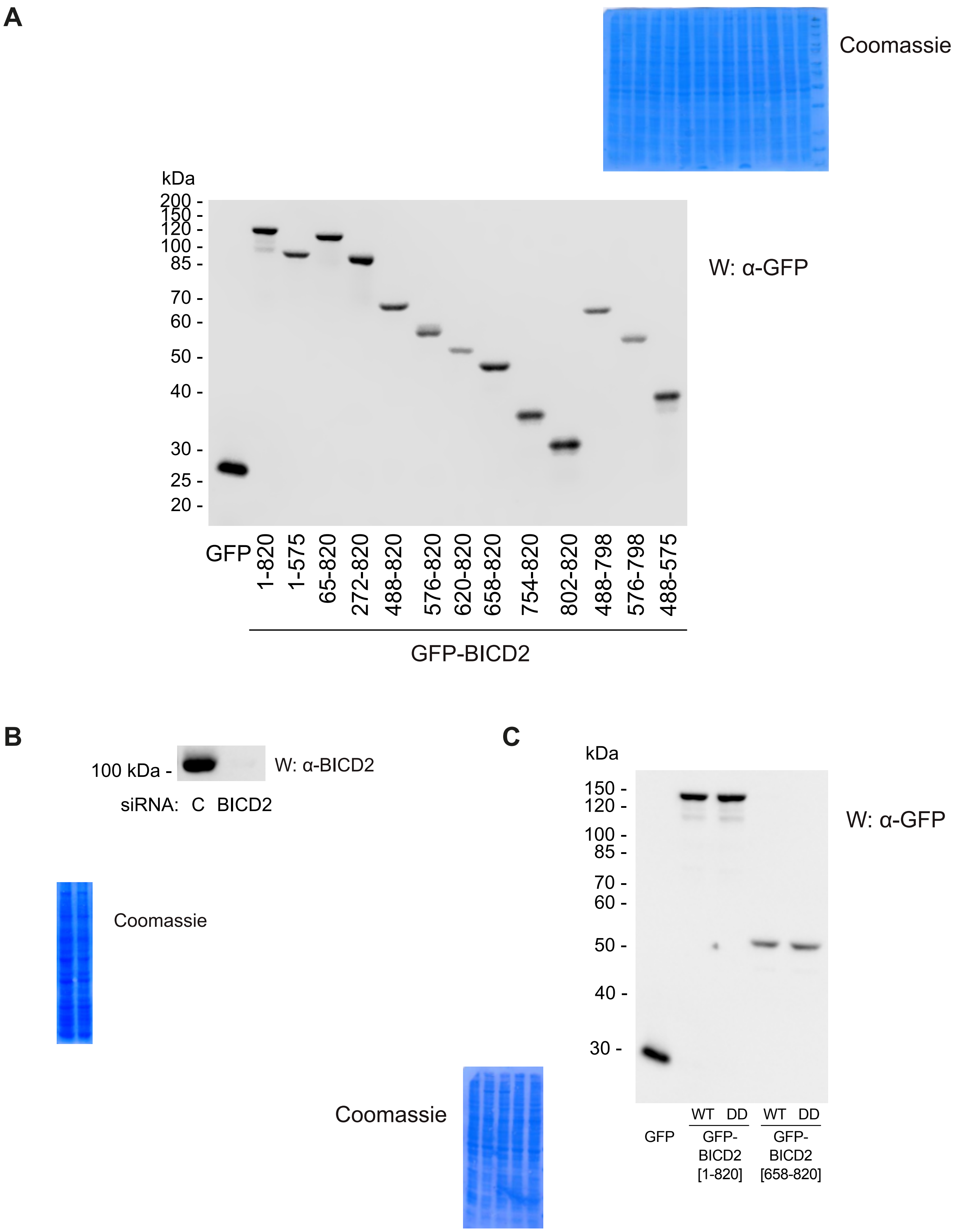
A
Coomassie
kDa
200 -
150 -
120 -
100 -
85 -
70 -
60 -
50 -
40 -
30 -
25 -
20 -
W: α-GFP
GFP
1-820
1-575
65-820
272-820
488-820
576-820
620-820
658-820
754-820
802-820
488-798
576-798
488-575
GFP-BICD2
B
100 kDa -
W: α-BICD2
siRNA: C BICD2
Coomassie
C
kDa
150 -
120 -
100 -
85 -
70 -
60 -
50 -
40 -
30 -
W: α-GFP
Coomassie
WT DD
WT DD
GFP
GFP-BICD2 [1-820]
GFP-BICD2 [658-820]
